# Epistatic evolution drives HLA-dependent CD8+ T Cell escape risk in diverse populations

**DOI:** 10.64898/2026.05.22.727291

**Authors:** David J. Hamelin, Jean-Christophe Grenier, Raphaël Poujol, Benoîte Bourdin, Bastien Paré, Shawn Simpson, Martin Smith, Hélène Decaluwe, Etienne Caron, Julie Hussin

## Abstract

Understanding how viral evolution shapes HLA-dependent T cell escape is crucial to identify individuals at risk of reduced cellular immunity to emerging variants. Nevertheless, we lack frameworks to model HLA diversity and the evolutionary feasibility of T cell–evading mutations. Here, we construct an HLA map capturing variation in epitope specificity across HLA-typed cohorts. Enhancing this framework with SARS-CoV-2 CD8⁺ T cell escape reveals heterogeneous escape across HLA-defined groups, with clusters enriched for HLA-B*07:02, HLA-A*03:01 and HLA-A*02:01 showing higher epitope loss. To assess the evolutionary plausibility of escape, we model viral sequence fitness using an epistasis-aware protein language approach trained on coronaviruses to systematically score mutations across viral lineages. We find that the fitness effect of mutations dynamically changes with evolving sequence context, and that T cell–evading mutations become fitter with additional escape mutations. This study links host HLA diversity to viral fitness landscapes for surveillance and vaccine design.

## MAIN

Pathogen-driven pandemics have repeatedly disrupted human societies, motivating extensive research on pathogen biology and public health interventions. Although not the deadliest, SARS-CoV-2 is the most comprehensively documented pandemic, genetically and epidemiologically, and thus essential for enhancing viral pandemic preparedness. To this end, understanding host genetic diversity and disease susceptibility is critical for dissecting host-pathogen interactions, pathogen evolution, and tailoring public health strategies.

The advent of personalized medicine has intensified the study of disease-genetic variant associations, many of which map to the highly polymorphic major histocompatibility complex (MHC) region^1^ . Within this region, the human leukocyte antigen (HLA) genes exhibit high levels of polymorphism and are central to adaptive immunity. Class I HLA molecules, expressed on all nucleated cells, present epitopes to CD8⁺ T cells, and their extensive allelic diversity (over 25,000 alleles across HLA-A, -B, and -C) underpins broad intra- and inter-individual protection against infectious diseases by enforcing distinct, allele-specific epitope-binding motifs^2–5^.

High-resolution HLA allele identification is required to capture population-scale HLA genetic diversity and can be performed using a variety of approaches, including imputation^6–9^, Next-Generation Sequencing (NGS) or Long-Read sequencing. Following HLA typing, a key step in studying HLA-disease associations is to define haplotype-based populations with distinct disease susceptibility. Population-genetic approaches using MHC Single Nucleotide Polymorphism (SNPs) and principal component analysis (PCA)^10–13^ tend to cluster individuals by ancestral lineage. While important for populational studies, these groupings may not always capture functional immunological features such as epitope specificity. Because the functional impact of HLA alleles is driven by specific amino acid residues in the peptide-binding groove, ancestry-based maps may obscure the true landscape of population-level immune vulnerability. An alternative is to use standardized HLA functional definitions such as 4-digit allele annotations, which capture amino acid differences relevant for epitope recognition^14–17^.

Rapid, affordable HLA typing and the identification of disease-susceptible HLA haplotypes are key for mitigating epidemics and guiding public health interventions. CD8+ T cells have been linked to milder COVID-19 symptoms and protection and may contribute to long-lived immunity^18–22^. However, SAR-CoV-2 has undergone significant diversification since its initial discovery, with multiple highly prevalent strains (Variants of Concerns, VoCs) responsible for its rapid spread. Both surveillance and evolutionary investigations have been greatly facilitated by genomic data-sharing platforms such as the Global Initiative on Sharing All Influenza data (GISAID)^23^. Most mutations were identified within the Spike Glycoprotein due its key role in immune evasion and cell host entry^24,25^ . Notably, SARS-CoV-2 diversification has generated mutations that escape T cell recognition^26–29^. Building on evidence that mutations can disproportionately disrupt immunogenic epitope presentation in an HLA-dependent manner ^30^, a growing body of evidence has showcased the accumulation of T cell evading mutations in several SARS-CoV-2 variants including Omicron (B.1.1.529), Omicron BA.1/2, and Omicron JN.1 ^26–29,31–42^. Notably, Omicron JN.1 has been characterized with the most extensive T cell escape to date. Overall, Omicron JN.1 was experimentally shown to lead to T cell escape in the context of 14 distinct CD8+ T cell epitopes across 9 HLA alleles, which is significantly more than previous lineages^33^ . Specific alleles such as HLA-A*24:02, A*02:01, B*27:05 and B*07:02 have been associated with CD8+ T cell escape and COVID-19 severity^43,44^, underscoring the need to map haplotype-specific susceptibility to T cell escape. While the strongest evidence of T cell escape are demonstrated using experimental approaches such as T cell assays, numerous predictive frameworks are available, such as NetMHCpan 4.1 for epitope-HLA predictions^45^, and PRIME 2.0 for HLA-epitope-TCR predictions^46^. These provide a high-throughput method for monitoring T cell escape across many HLA alleles, antigens, and viral strains.

Beyond surveillance of viral immune escape, forecasting viral evolution requires understanding how mutations affect viral fitness. Epistatic interactions between mutations, where the effect of a mutation depends on the surrounding sequence context, have been shown to play key roles in several aspects of viral evolution. In the context of immune escape, epistasis was shown to lower the barrier to antibody escape in SARS-CoV-2 variants^47–49^, underscoring the importance of evaluating mutation fitness within lineage-specific genetic backgrounds. High-throughput functional assays such as Deep Mutational Scans (DMS) have enabled large-scale interrogation of viral mutations but remain labor-intensive and limited in their ability to probe the combinatorial effect of mutations across diverse sequence contexts. To mitigate this, recent advances in computational methods enable scalable estimation of mutation fitness, both individually and against vast arrays of sequence backgrounds^50^ . Importantly, the predictability of deep learning models toward a virus of interest can be increased by learning from sequences from its viral family^51–53^. Protein Language Models (PLM), such as Evolutionary-Scale Modeling 2 (ESM-2), a transformer trained on 65 million UniRef protein sequences, can provide a fitness proxy for a mutation by masking its position and evaluating the model’s likelihood for that residue^51,54^.

In this work, outlined in **Fig 1**, we introduce a framework that groups individuals by class I HLA haplotype, emphasizing shared epitope preferences. Across multiple cohorts (The 1000 Genomes Project^10^ , UK Biobank^55^ , and the RECOVER cohort^56,57^, an Oxford Nanopore-based HLA-typed cohort), these functional clusters reveal structure that is not captured by classical population genetic approaches and set the stage to identify populations with elevated susceptibility to CD8+ T cell escape. We then leverage both predicted and experimentally validated SARS-CoV-2 CD8+ T cell escape mutations across these cohorts, identifying HLA-defined populations vulnerable to epitope loss. Finally, we assess the evolutionary plausibility of T cell escape using an epistasis-aware PLM. We identify T cell escape mutations that increase in fitness throughout the pandemic and as their respective viral strains accumulate additional T cell escape substitutions. Together, these analyses unify host HLA landscapes with viral fitness landscapes to characterize both immunologically consequential and evolutionarily accessible T cell escape, outlining a strategy to guide variant surveillance and next-generation vaccine design.

**Figure 1.**
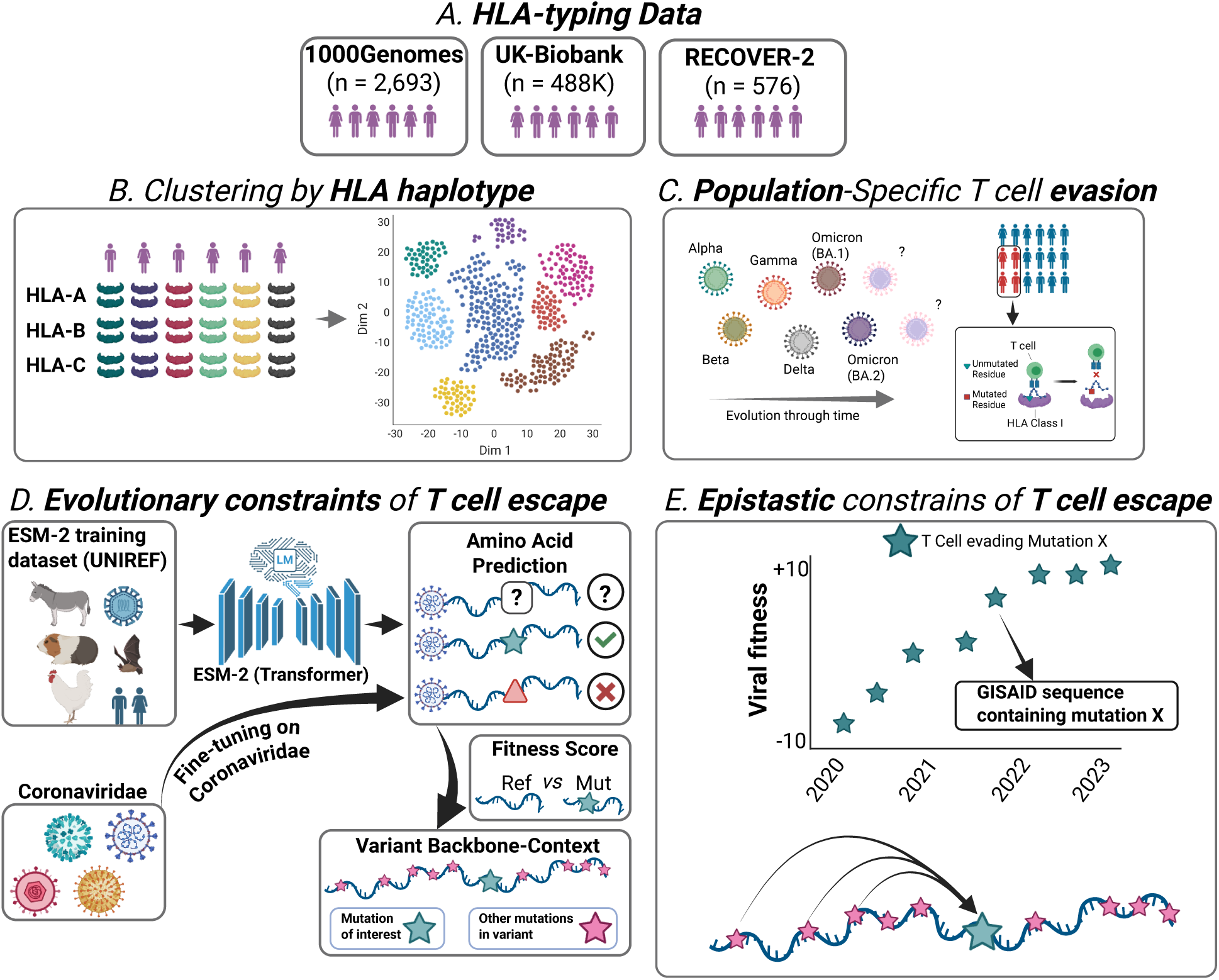
Graphical description of the study. ***(A)*** Classical HLA class I profiles were acquired from several cohorts. ***(B)*** Individuals were clustered together based on the functional similarity in their HLA alleles. ***(C)*** HLA profile-defined populations at risk of T cell evasion to SARS-CoV-2 variants were identified using a combination of epitope predictions as well as peer-reviewed T cell evasion data. ***(D)*** The evolutionary constraints of T cell evading mutations were interrogated using ESM-2, a Protein Language model, which we fine-tuned on *Coronaviridae* sequences. ***(E)*** The evolutionary constraints of mutations were further described by investigating epistatic dependencies between mutations and determining the impact of evolving sequences on the fitness of T cell evading mutations

## RESULTS

### Immunogenetic structure differs fundamentally from ancestry-defined population groups

Understanding how HLA class I diversity is structured across human populations is essential for predicting differential susceptibility to T cell escape. To this end, we sought to construct an immunologically meaningful population map to characterize population-level vulnerability to T cell escape. Standard population stratification methods typically rely on genomic ancestry, often obscuring functional similarities in immune potential across diverse individuals. To address this limitation, we developed HLAScape, a novel computational workflow that reorganizes populations into distinct HLA haplotype–defined clusters based on functional resemblance. By adapting the Jaccard index to quantify HLA haplotype overlap and combining t-SNE dimensionality reduction (Methods), we generated an embedding in which neighboring individuals share similar 6-allele HLA class I haplotypes, providing a proxy for similarity in antigen presentation potential.

This approach was applied to the 1000 Genome Project (1KGP) to investigate HLA haplotypes across genetically diverse populations (**Fig 2A, B**). We identified 11 distinct HLA-defined clusters (**Fig 2A**) and visualizing the top four alleles across HLA-defined clusters puts in evidence intra-cluster overlap and inter-cluster diversity (**Fig 2B**). To assess clustering success, we developed a K-nearest neighbour (KNN)-based interpretability metric (Methods, Supp Notes 1.1), to determine that individuals shared on average 3.21 of 6 HLA class I alleles with their nearest clustering neighbours (Global KNN_HLA_ = 3.21), effectively more than expected under label shuffling (Global KNN_HLA_ = 0.57) **(Fig 2C)**. In addition, the resulting HLA map partially captured populational structure with individuals sharing ancestry labels with 55% of nearest neighbors (Global KNN_ancestry_ = 0.55, see Methods) **(Ext. Data Fig 1A)**. Nevertheless, the distribution of ancestry labels across clusters suggests the clustering to be HLA haplotype-driven more than ancestry-driven **(Ext. Data Fig 1C)**.

**Figure 2.**
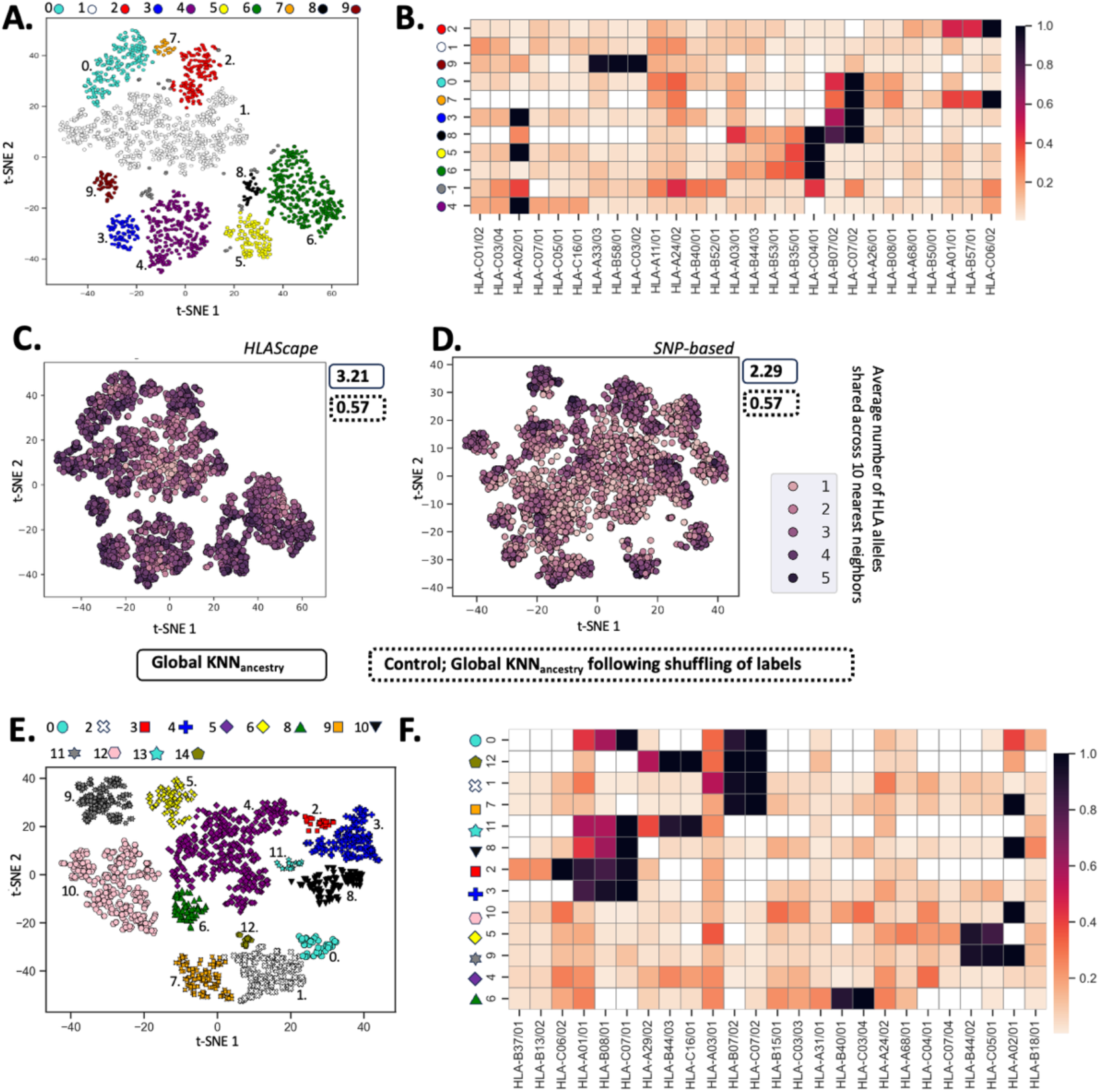
A novel strategy to interrogate HLA haplotype diversity offers a unique perspective on HLA haplotype populational structure. **A.** Clustering of 1KGP individuals based on HLA profiles. Dots correspond to unique HLA profiles (HLA-A, B and C), clustered by HLA-allele similarity. **B.** Top HLA alleles are shown for each cluster. Only the top 4 alleles per cluster are shown **C.** Global KNN_HLA_ interpretability metric for HLA haplotype clustering, measuring the average number of shared alleles across the 10 nearest neighbors for a given HLA profile. Global KNN_HLA_ (*) is the average number of shared HLA alleles (out of 6) among the 10 nearest neighbors, across the cohort; control (**) is the Global KNN_HLA_ following shuffling of labels. **D.** SNPs were extracted from the MHC region in the 1KGP (MAF > 1%, LD R² < 2 in 50-SNP blocks, n = 467), then clustered using t-SNE on the first 20 Principal Components from SNP genotypes. **E.** Clustering of UKB individuals based on HLA profiles (same as A). **F**. Top HLA alleles are shown for each cluster (same as B).

The proposed framework was contrasted with a classical ancestry-driven population genetics approach, consisting of clustering individuals using SNP genotypes in the MHC region with PCA (Supp Notes 1.2.1). This strategy reflected neutral ancestry differences (**Ext. Data Fig 2**) while yielding higher ancestry clustering (Global KNN_ancestry_ = 0.70) and lower clustering by HLA haplotype similarity (Global KNN_HLA_ = 2.29) than HLAScape **(Fig 2D, Ext. Data Fig 1B,D)**. As an intermediate, leveraging SNP restricted to peptide-presentation genes (HLA-A/B/C) partially recovered HLA haplotype clustering (Global KNN_HLA_ = 2.76) at the cost of ancestry-based clustering (Global KNN_ancestry_ = 0.49) **(Ext. Data Fig S1, Fig S1)**. Hence, HLA allele definitions capture variation that is more directly relevant to antigen presentation than neutral SNP variation and provides a functionally informed view of peptide-binding diversity.

To determine whether the clustering patterns generated by HLAScape were method-specific, we explored additional alternative dimensionality reduction and pre-processing strategies (**Fig S2,** Supp Notes 1.2.2). Notably, Multi-Dimentional Scanning (MDS) resulted in inferior HLA haplotype clustering (Global KNN_HLA_ = 1.75). Nevertheless, this yielded clusters with HLA compositions highly similar to those from t-SNE, indicating that the observed structure does not depend on the dimensionality reduction technique used (**Ext. Data Fig 3, Fig S3**, Supp Notes 2.2.3.)

To evaluate the robustness of the structure and the ability of this approach to scale to larger cohorts, we applied our framework to a second cohort, the UKB (**Fig 2E, F**), which is primarily composed of individuals of European ancestry, and contains a broad set of unique class I HLA allele combinations (n = 94,225 HLA subtype A, B, C combinations). UKB yielded 13 distinct HLA-defined clusters (**Fig 2F**), recapitulating patterns observed in the European subset of 1KGP (**Fig S4**). Finally, to ensure applicability in a disease-focused setting, we mapped 576 SARS-CoV-2 convalescent individuals from the RECOVER cohort into an HLA haplotype landscape using high-throughput Oxford Nanopore-based HLA typing (Methods, Supp Notes 2.1) (**Fig 3A, B**). RECOVER individuals distributed evenly across the existing cluster structure observed for the UKB (**Fig 3C**) and formed seven coherent subgroups (**Fig 3B-E**) with HLA structure similar to that the European population of 1KGP **(Fig S4A)**, confirming that the HLA-defined axes of variation identified in population cohorts are preserved in this clinical cohort. Together, these analyses demonstrate that immunogenetic structure is robust and reproducible across cohorts, establishing a unified HLA haplotype framework for assessing CD8⁺ T-cell escape risk.

**Figure 3.**
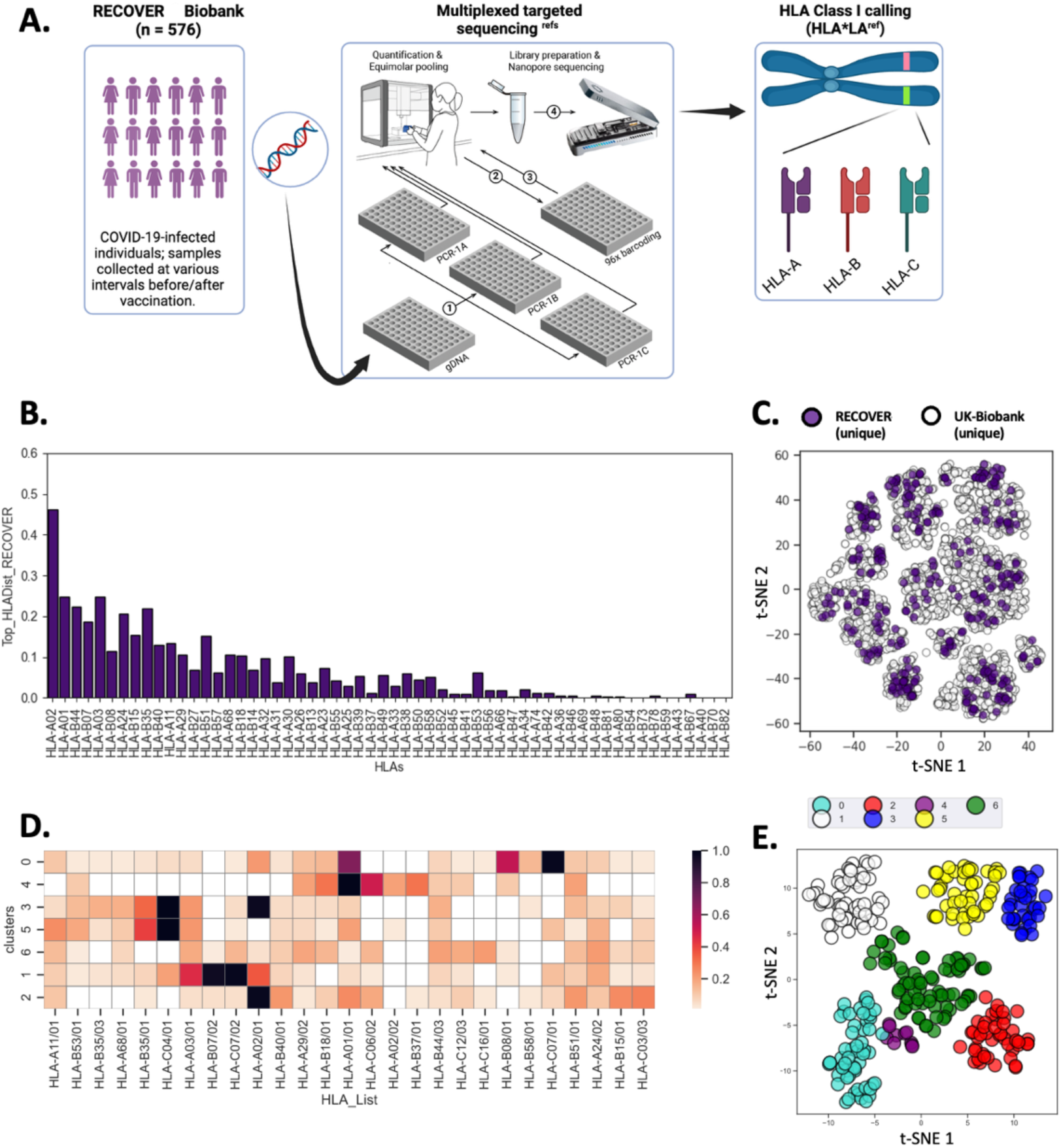
Population-scale HLA-typing enabled by an affordable, high throughput Oxford Nanopore-based approach. **A**. Population-scale HLA typing. Briefly, PBMCs were acquired from SARS-CoV-2-convalescent patients (left). Targeted HLA amplification and sequencing was performed according to Stockton *et al,* 2020, and sample multiplexing was conducted as per the Oxford Nanopore Rapid Barcoding Kit 96 protocol (center). Finally, HLA class I calling was done with the HLA*LA software (Dilthey *et al*., 2019) (right). **B.** Distributions of HLA alleles in the RECOVER cohorts. **C.** Clustering of UKB and RECOVER cohorts HLA profiles (see methods). **D.** Top 6 HLA alleles are shown across all cluster (colored boxes). **E.** Clustering of RECOVER HLA profiles, colored based on clustering using DBSCAN.

### HLA-driven immunogenetic clusters reveal population variation in CD8⁺ T-cell escape risk

We sought to evaluate if HLA-defined immunogenetic clusters differ systematically in CD8⁺ T cell escape risk across populations (**Ext. Data Fig 4**). T-cell evasion across populations was analyzed using two distinct approaches: (i) by considering experimentally validated T-cell escape only, and (ii) by predicting T-cell evasion *in silico* (Methods). The resulting haplotype-specific escape burden was then used to characterize inter-cluster variability.

HLA-dependent T-cell escape by SARS-CoV-2 point mutations was initially assessed across the haplotypes in the context of the variant Omicron JN.1 due to the variant’s extensive demonstrated T-cell evasion^33^. Omicron JN.1 mutations lead to T-cell escape across multiple clusters for both experimentally validated **(Fig 4A, Ext. Data Fig 5**) and for predicted evasion (**Ext. Data Fig 6**). In all interrogated cohorts, experimentally validated T-cell escape was particularly elevated in clusters enriched in HLA alleles B*07:02, corroborating prior findings^30^, as well as HLA allele A*02:01 and A*24:02 **(Fig 4A**). This escape pattern was further emphasized when considering the immunogenicity-weighted escape score (**Ext. Data Fig 5**). Notable escape events (non-exhaustive) include the mutational hotspots, N450D/L452W/L455S and S371F/S373P/S375F/T376A shown to abrogate the HLA-A*24:02 T cell epitopes NYNYLYRLF (NY**D**Y**W**YR**S**F) and LYNSASFSTF (LYN**F**A**P**F**FA**F); in the mutation Q229K which leads to the abrogation of three distinct A02:01 epitopes; and in the mutation P681R/H which leads to the loss of the SPRRARSVA epitope. Looking at other variants beyond Omicron JN.1, we see that T-cell escape patterns across HLA haplotype groups varied greatly throughout the pandemic, with earlier variants leading to lesser T-cell escape **(Fig 4B-C, Ext. Data Fig 5, 6**). Importantly, the set of affected haplotype groups appears variant dependant, although Omicron JN.1 leads to extensive T-cell escape across the greatest number of distinct clusters. Predicted T-cell escape partially recapitulated experimentally validated escape events but also suggested broader multi-cluster escape across cohorts and variants, pointing to additional candidate escape mutations (**Ext. Data Fig 6**). To determine whether the observed escape patterns could arise by chance, we generated *in silico* SARS-CoV-2 variants in which mutations were randomized within known Omicron mutational hotspots (Supp Notes 2.4). While a subset of synthetic variants (25%) exhibited greater predicted escape potential than Omicron BA.2, the observed escape profiles fell well above random expectations **(Fig S5)**, indicating a structured T-cell escape landscape. Together, these results show that T-cell escape is highly heterogeneous, with specific groups of HLA haplotypes susceptible to greater T-cell escape than others.

**Figure 4.**
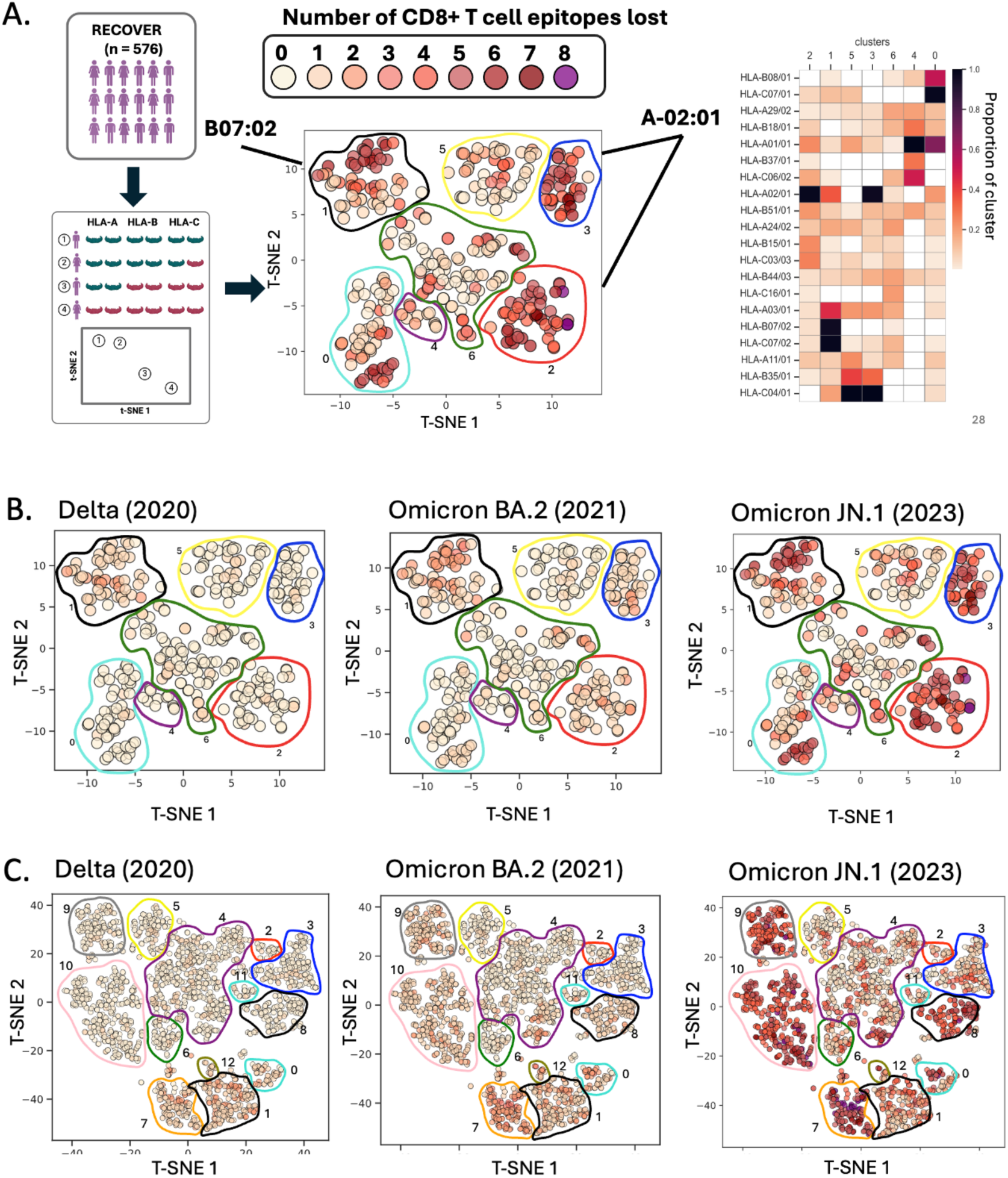
Characterization of CD8+ T cell epitope loss across multiple cohorts. **A**. Illustrative example of population-specific evasion across the RECOVER cohort. All epitopes, their corresponding variant-specific mutations, and the resulting escape event considered here were acquired from the literature (see Methods). Total T cell evasion per participant was determined by quantifying the number of epitopes lost to variant mutations based on the complete participant HLA class I HLA profile (shown on the right). **B**. T cell evasion across the RECOVER cohort by mutations specific to Delta, BA2, and JN1. **C**. T cell evasion across the UKB by mutations specific to Delta, BA2, and JN1

T-cell escape patterns across HLA haplotype groups were highly reproducible across the UKB, RECOVER, and the European subset of the 1KGP (**Fig S6**), indicating robust immunogenetic structure within populations of similar HLA composition. By contrast, African, East Asian, and South Asian 1KGP populations exhibited highly distinct HLA clustering and T-cell escape profiles reflecting differences in HLA allele frequencies (**Fig S7**). This highlights the importance of both the global HLA diversity and ancestry-dependent immunogenetic structure in shaping population-level vulnerability to T-cell escape. Therefore, host immunogenetics susceptibility is key to preparedness for emerging variants, while lineage-specific viral evolution will constrain what escape mutations can arise.

### Leveraging an epistasis-aware Protein Language Model to forecast T-cell evading mutations

To anticipate future T-cell escape under evolutionary constraints, we developed a protein language model (PLM) framework to estimate the context-dependent fitness of escape mutations across viral backgrounds (Methods). Because viral mutations act within a co-evolving genome, the fitness of a single mutation is treated as a dynamic value that depends on the surrounding sequence context, as opposed to a fixed, inherent value. Due to context learning, PLMs are well-suited to implicitly capture epistatic relationships between a mutation of interest and any strain-specific genetic background. To this end, we fine-tuned the ESM-2 PLM on Coronaviridae sequences (ESM2-CoV) and used it to interrogate fitness effects of SARS-CoV-2 Spike mutations.

To test the concordance between ESM2-CoV-estimated mutation fitness and their observed pandemic success, we performed *in silico* mutagenesis analysis by computing the fitness of all Spike point mutations and compared these values to their frequencies in 50,000 subsampled SARS-CoV-2 sequences from GISAID. Viral fitness was only predicted to be elevated for mutations that were also frequently observed throughout the pandemic, suggesting a concordance between predicted and observed mutation fitness (**Fig S8A**, Supp Notes 3.1). To evaluate variant-dependent fitness, we used Spike DMS data from Bloom et al. (BA.2 and XBB.1.5 backbones) and found that high DMS scores only corresponded to high ESM2-CoV scores when mutations were tested against their native backbones (**Fig S8B-D**, Supp Notes 3*.1*.). We performed additional checks to validate if our method can disentangle the contribution of mutations while considering the viral backgrounds. *In silico,* we introduced unobserved, low-fitness mutations into Spike sequence backgrounds sampled across the pandemic to test whether model inference is disproportionately driven by sequence context (**Ext. Data Fig 7D**). These mutations were given exceedingly low fitness scores in nearly all backgrounds, demonstrating that background alone is insufficient to result in high inferred fitness.

Leveraging our framework, we tracked mutation fitness over time across large numbers of viral sequences using GISAID (Supp Notes 3*.2*). For each mutation, we collected all sequences carrying that change and estimated its fitness in the native Spike sequence context in which it was observed. As illustrative examples, Spike N460K and Spike G446S, both highly prevalent mutations present in multiple VoCs, showed gradual increases in fitness as surrounding mutations accumulated, highlighting strong background dependence (**Ext. Data Fig 7A-B**). Spike mutations from Delta, Omicron BA.2 and Omicron JN.1 revealed mean fitness values spanning slightly deleterious/neutral to beneficial (**Ext. Data Fig 7C; S9A–B**). To gain insight on variant-specific epistatic structure, we estimated the fitness of each variant-defining mutation in the presence of every other single mutation from the same VoC **(Fig 5)**. This revealed paired mutation interaction schemes distributed across the Spike protein and enriched in and near the RBD. The strongest interactions preferentially involved proximal sites, consistent with shared structural or functional constraints. Comparable interaction patterns were observed in analyses of mutation triplets (i.e., a query mutation with two companion mutations) (**Fig S10,** Supp Notes 3.3).

**Figure 5.**
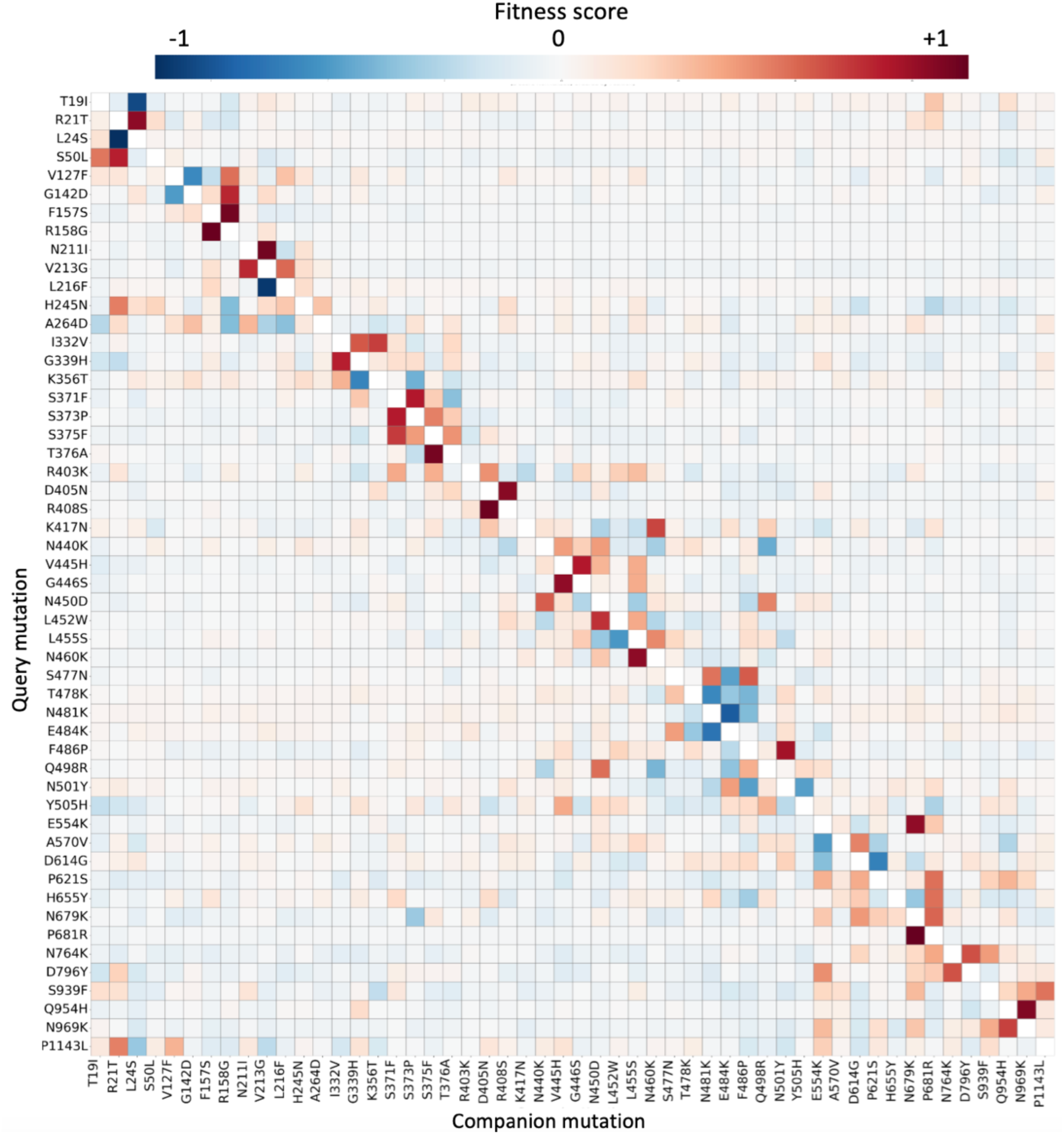
Omicron JN.1 two-mutation interaction map. The fitness of each Omicron JN.1 mutation was estimated in the presence of a single other JN.1 mutation for all JN.1 mutations to create a two-mutation interaction map. Mutations are ordered by mutation position along the Spike protein. Here, the y-axis corresponds the the “query mutation” (for which the fitness was assessed), while the x-axis corresponds to the “companion mutation” which was added to the backbone as a single companion mutation. Since the aim was to visualize the distribution of fitness effects of each query mutation across all companion mutations (and not to compare query mutations to each other), all fitness metrics associated with one query mutation was normalized using a z-score ([data value – average of all points]/standard deviation).

Ultimately, we used this framework to investigate T-cell evading mutations. Specifically, we examined how viral sequences evolved alongside the accumulation of escape mutations, assessing the relationship between viral fitness and T-cell escape. We prioritized experimentally validated T-cell evading mutations from the Omicron JN.1 subvariant, which were largely found to increase in fitness along with evolving sequence backgrounds (**Fig 6A**). Notably, this increase in fitness was accompanied by an increase in co-occurring T-cell escape mutations and affected HLA alleles. For instance, the Spike mutation P681R—which mediates T-cell escape of the immunogenic peptide SPRRARSVA (HLA-B*07:02)—shows increased fitness in later variant backbones and co-occurs with a growing set of T-cell evasion substitutions, alongside an expanding impact across HLA alleles **(Fig 6A)**. A similar trend was observed in the Spike mutation S371F, involved in the T cell escape of the peptide LYNSASFSTF (HLA-A*24:02) **(Fig 6A)**, again linking improved evolutionary accessibility with progressive broadening of T-cell escape.

**Figure 6.**
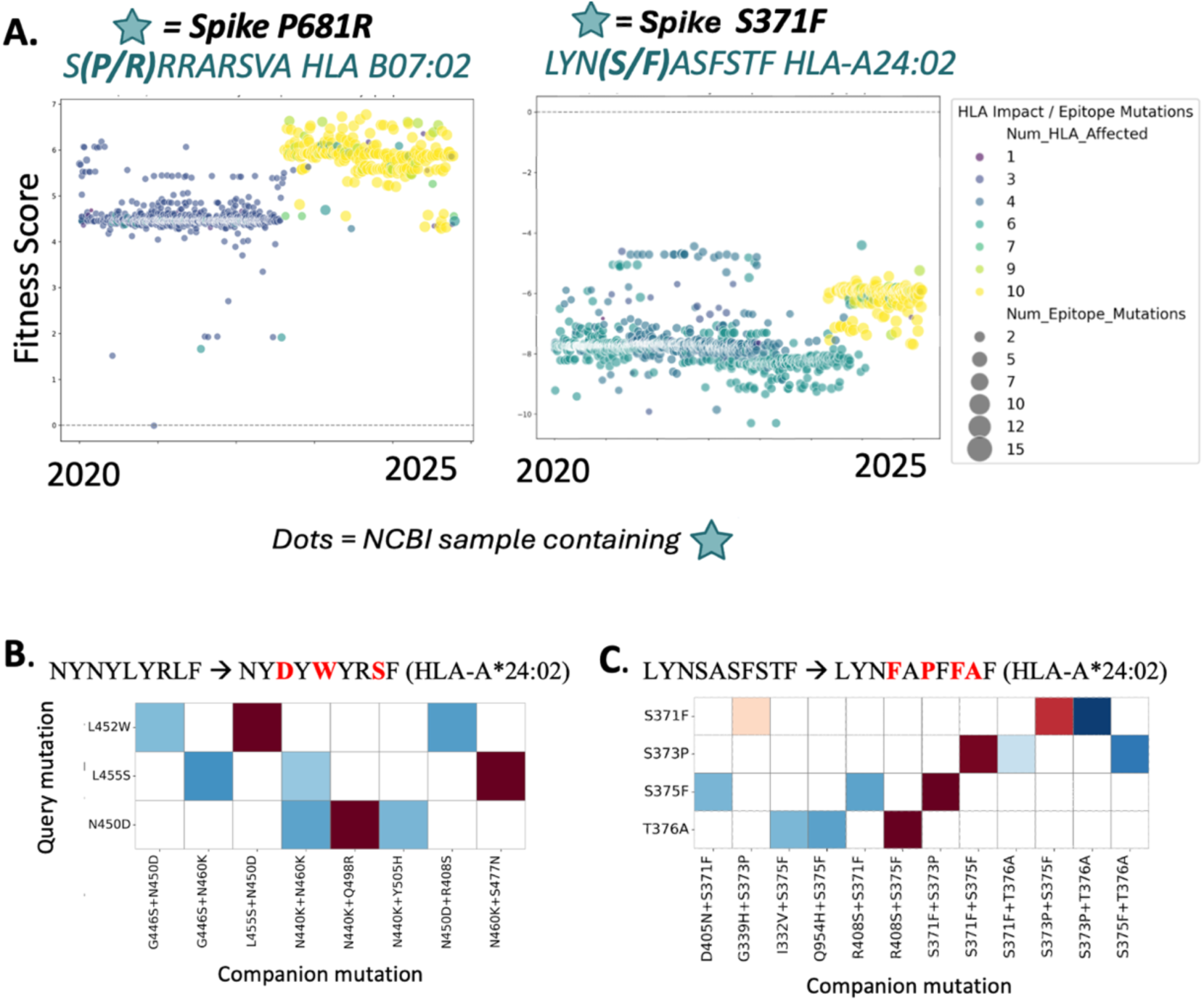
**(A)** Estimated fitness of known T cell evading mutations in native Spike protein background spanning the pandemic. Dot color and size correspond to the quantity of other T cell evading mutations in native sequences, with dot color representing the number of HLA alleles impacted and the dot size corresponding to the total number of epitopes affected **(B-C)** Three-mutation interaction map for query mutations found within t-cell evading hot-spots. The fitness of each mutation from two T cell evading hotspots (NYNYLYRLF è NY**D**Y**W**YR**S**F **(B)** and LYNSASFSTF è LYN**F**A**P**F**FA**F **(C)**) was estimated in the presence of two other JN.1 mutation for possible JN.1/T cell hot spot 3-mutation combinations to create a three-mutation interaction map. In all figures, the y-axis corresponds the “query mutation” (for which the fitness was assessed), while the x-axis corresponds to the “companion mutation(s)” added to the backbone as a single companion mutation (A) or double companion mutations (B, C). Since the aim was to visualize the distribution of fitness effects of each query mutation across all companion mutations (and not to compare query mutations to each other), all fitness metrics associated with one query mutation was normalized using a z-score ((data value – average of all points)/standard deviation).

Consistent with the increasing breadth of affected HLA alleles, higher PLM-inferred viral fitness of T-cell-evading mutations was associated with broader escape across HLA-typed cohorts. The highest-fitness backgrounds showed the strongest escape effects across HLA clusters (**Fig 6A**). Of particular interest are two mutational hotspots that were both experimentally associated with the escape of immunodominant epitopes. The N450D/L452W/L455S JN.1 mutational hotpot and the S371F/S373P/S375F/T376A BA.2/JN.1 mutational hotspot were shown to abrogate the HLA-A*24:02 T cell epitopes NYNYLYRLF (NY**D**Y**W**YR**S**F) and LYNSASFSTF (LYN**F**A**P**F**FA**F), respectively (Supp Notes 3.4)^31,33^. These represent putative examples of viral adaptations to host T-cell immunity, due to the hypermutation of both epitopes as well as the prevalence of their restricting HLA allele. All mutations within both hotspots were found to have either neutral or beneficial fitness effects on the virus when assessed within their respective contexts (**Ext. Data Fig 7C** addition, interaction analyses focused on these two hotspots revealed strong background dependence, with many substitutions only tolerated or becoming advantageous in the presence of intra-hotspot and hotspot-proximal mutation partners (**Fig 6B-C**). To explore the full range of potential epistatic relationships associated with these hotspots, we assessed the fitness of each hotspot mutation in the presence of all possible point mutations in a 40-position window **(Ext. Data Fig 8, S11-S12)**. Alongside recapitulating the beneficial effect of known hotspot mutations on the additional mutations of interest, we identify additional non-observed high-fitness mutation pairs, thus suggesting potential routes for further expansion of these hotspots. Together, these results suggest that SARS-CoV-2 mutational trajectories are not only leading to an increase in viral fitness, but to an increase in T-cell escape at immunodominant epitopes restricted by common HLA alleles, thereby expanding both the magnitude and the population breadth of T-cell escape.

## DISCUSSION

The COVID-19 pandemic has profoundly impacted economies, social norms, and public health, driving unprecedented international collaboration, open science, and data-sharing. It has also underscored the need for preparedness strategies for future pandemics and SARS-CoV-2 outbreaks, including workflows to assess T-cell escape across genetically diverse populations. However, the global diversity of HLA alleles means that the impact of VoCs on T-cell immunity remains incompletely understood. In this work, we develop and visualize a functional map of human HLA class I haplotype diversity to identify populations at risk of T-cell escape, and establish an epistasis-aware protein language modelling framework to investigate the fitness dynamics of immune-evading variants, enabling integrated prediction of viral sequence fitness.

Our HLA map groups individuals according to similarities in their HLA class I haplotypes, as defined by shared allele composition and associated epitope binding preferences. In the resulting embedding, proximity between individuals reflects similarity in antigen presentation potential, whereas classical population genetics approaches only partially capture this through ancestry structure. We introduced and applied the HLAScape approach across multiple cohorts of varying sizes, demonstrating scalability to large datasets (e.g., UKB n=488,221 HLA-imputed individuals presenting 94,225 unique HLA class I profiles). This highlights the importance of representing global HLA diversity in genomic surveillance efforts, as reliance on ancestry-based groupings risks overlooking immunogenetically vulnerable populations that are underrepresented in current biobanks. Continuous monitoring of T-cell escape in ongoing and future pandemics will require renewed emphasis on HLA-typing campaigns across emerging databases, biobanks, and clinical cohorts. The high cost of standard HLA-typing methods has remained a barrier to large-scale efforts. Here, our high-throughput long-read HLA-typing approach, enabled by laboratory automation and scalable multiplexing, allowed cost-effective profiling of a clinical cohort and the application of our framework at scale. This strategy may facilitate future integration of host immunogenetic diversity with viral evolution in surveillance and vaccine design, including the potential for population-scale or personalized vaccination strategies informed by anticipated immune escape. Notably, analysis of the RECOVER cohort was consistent with results from the UKB and the European subset of the 1KGP population, supporting the replicability of our HLA haplotype visualization across cohorts.

Overall, our approach organizes individuals into HLA-defined clusters that provide a framework for quantifying and comparing T-cell escape risk across populations. Overlaying SARS-CoV-2 variation onto this HLA map reveals marked heterogeneity in how viral evolution impacts different HLA-defined groups. By integrating predicted and experimentally validated escape mutations, we identify multiple haplotype clusters with elevated susceptibility to T-cell escape, particularly those enriched for alleles such as HLA-A*02:01, HLA-A*24:02, and HLA-B*07:02. Importantly, patterns of population-specific immune escape vary across viral variants, with Omicron JN.1 exhibiting the most extensive escape across haplotype clusters, consistent with prior observations^33^.

To complement our characterization of T-cell escape across a genetically diverse host population, we examined the evolutionary constraints governing escape mutations to anticipate emergence and identify at-risk populations early. To do so, we developed a deep learning model fine-tuned on Coronaviridae, ESM2-CoV, to estimate viral fitness at amino-acid resolution. By learning higher-order dependencies among residues, PLMs inherently incorporate epistasis and can be applied across diverse mutational backgrounds. Accordingly, we treat mutation fitness as dynamic, varying with sequence context evolves, rather than static. We show that ESM2-CoV captures this evolving fitness landscape, with the same mutation exhibiting markedly different effects across strains and over time, including shifts from detrimental or neutral in early variants to advantageous in later variants. Finally, we find that most interactions are constrained to local epistatic windows, with occasional distal effects, providing a framework to interrogate interdependent mutational hotspots in emerging Omicron sublineages.

Applying this approach to T-cell escape mutations, we quantified the number of co-occurring - escape mutations within individual viral sequences across the COVID pandemic (GISAID, 2020-2025). We observe an increase in the context-dependant fitness of T-cell evading mutations, accompanied by the accumulation of escape mutations and an expanding set of affected HLA alleles within viral strains. Our in-depth investigation of two mutational hotspots found within immunodominant T-cell epitopes shows a strong evolutionary interdependence among mutations within each hotspot. Together, these observations support a model in which SARS-CoV-2 gradually explores immune-escape trajectories that were initially constrained by fitness costs but become more accessible as compensatory mutations accumulate. Although demonstrated here for SARS-CoV-2, this framework could extend easily to other viruses with T-cell-mediated immunity, pathogens under strong CD8+ T-cell selection, and emerging viruses in future pandemics.

This work has several limitations. While our HLA clustering strategy considers a binary similarity metric between individual HLA alleles, we could further develop this by considering partial or graded similarity. This could provide a complementary view of the data, while incorporating additional fine-grained functional information such as predicted peptide-binding profiles. Our immunogenetic analyses focus on class I HLA–restricted CD8+ T-cell escape and do not capture other immune axes such as antibody-mediated selection, CD4+ T-cell responses, or innate immunity. However, our protein language modelling framework is agnostic to immune mechanism and captures general viral fitness constraints and, thus, offers a flexible foundation for future investigation of multiple selective pressures. Because UKB and RECOVER cohorts are enriched for European ancestry, population-specific immune escape in non-European populations was less thoroughly examined, although we observe distinct immunogenetic structures and escape profiles in non-European populations from the 1KG cohort. For all viral sequence analyses, we focused on the Spike protein as it provides a biologically and clinically relevant test case, but the framework itself is readily extensible to other SARS-CoV-2 proteins and other viruses. Both experimentally validated and predicted T-cell escape events are constrained: validation assays focus on common HLA alleles, omitting rare but possibly relevant ones, while *in silico* predictions are subject to high false-positive rates, especially for HLA alleles underrepresented in training data. Finally, PLM-based fitness scores remain proxies and should be interpreted as a constraint on variant emergence rather than direct measures of replication or transmission success.

In summary, our study outlines an adaptable and scalable strategy that integrates host immunogenetic structure with viral fitness constraints to identify populations at elevated risk and prioritizing mutations that are both immunologically consequential and evolutionarily plausible. We lay the groundwork for forecasting T-cell evading events, while also providing key insights for prioritizing emerging mutations for experimental validation and for guiding the monitoring of epitopes relevant to common HLA haplotypes. This combined host-virus framework supports the design of next-generation vaccines robust across genetically diverse populations, while being broadly applicable to other rapidly evolving pathogens under T-cell-mediated immune pressure.

## ONLINE METHODS

### Cohorts and HLA data sources

#### HLA data for UK Biobank (UKB) and 1000 Genome Project (1KGP)

HLA typing data was acquired from both the 1KGP (n = 2693) and UKB (n = 488,221). 1KGP HLA alleles are publicly available and were typed by performing in silico typing using PolyPheMe (with an IMGT/HLA reference database) on 1KGP exome sequence^58^ . UKB classical HLA alleles were imputed using HLA*IMP:02, using UKB genotype calls and a multi-population reference panel^7,59^ . For both cohorts, we obtained four-digit (two-field) HLA alleles at HLA-A, HLA-B and HLA-C. Allele nomenclature was harmonized across cohorts to a consistent two-field resolution prior to downstream analyses. For the 1KGP, we investigated HLA genetic diversity across cohort participants. Briefly, extracting SNPs from the complete MHC region (8,574 SNPs) and HLA A, B, C, (467 SNPs). For control, SNPs were randomly selected across chr6 (excl. MHC region) with SNP counts matching those corresponding to the full MHC region and HLA A, B, C. SNPs were extracted using plink v2^60^ , and filters for minor allele frequency (MAF) and linkage disequilibrium (LD) were applied (MAF >= 1%; LD R^2^ < 2 in blocks of 50 SNPs). In all cases, SNPs were clustered using t-SNE on the first 20 Principal Components from SNP genotypes.

#### RECOVER subjects and sample collection

The study subjects were composed of previously infected health care workers (HCWs) who were recruited following a PCR-confirmed SARS-CoV-2 infection as part of the RECOVER study (n=576). Blood samples were collected at enrollment around 6.1 ± 2.4 months after infection into acid-citrate-dextrose tubes (ACD, BD) in each of the five participating centers in the Province of Québec, shipped to the Mother-Child Biobank at the CHU Sainte-Justine where peripheral blood mononuclear cells (PBMCs) were isolated according to standard operation procedures (SOPs) using SepMateTM tubes (Stemcell Technologies, Canada). PBMCs were cryopreserved in complete RPMI (Gibco) with 10% DMSO and stored in liquid nitrogen until used. Participants were recruited from August 17, 2020, to April 8, 2021. High-throughput HLA-typing of the RECOVER cohort was performed using a long-read sequencing approach (Supp Notes 2.1). Briefly, targeted amplification of HLA class I loci was adapted from Stockton et al, 2020^61^ for 96-well plate processing and laboratory automation. Amplicons were quantified, normalized, and pooled prior to library preparation using a combination of Oxford Nanopore Technologies ligation-based (SQK-LSK109 with EXP-NBD196) and rapid (SQK-RBK111.96) barcoding protocols. Sequencing was performed on MinION (R9.4.1), and HLA genotypes were inferred using HLA*LA^62^ (Supplementary File 1).

### HLAScape: HLA haplotype representation and immunogenetic clustering

#### Intra- and inter-cohort HLA class I-based diversity

The HLA class 1-based diversity within the cohorts interrogated in this study was determined by considering all 6 HLA alleles identified for each participant. To do so, the Jaccard distance was applied^63^ . Briefly, the Jaccard index was implemented to reflect the similarity in HLA profiles between participants, as shown below:

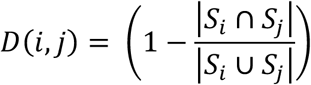

Here, the similarity in HLA class I profiles between participants *i* and *j*, described by the Jaccard distance D(*i, j*), is determined by computing the number of shared HLA alleles between both participants,|𝑆*_i_* ∩ 𝑆*_j_*|, as well as the total number of unique HLA alleles found amongst both participants ,|𝑆*_i_* ∪ 𝑆*_j_* | Subsequently, an HLA profile diversity map was constructed by converting the Jaccard distance matrix to 2-dimentional space using t-distributed stochastic neighbor embedding (t-SNE)^64^ . Finally, the clusters generated by t-SNE were identified using the density-based clustering algorithm DBSCAN^65^.

#### KNN-based interpretability metric for assessing clustering

A K-nearest-neighbor (KNN)-based interpretability metric was established to assess the degree of success with which individuals were clustered, and to compare the impact of varying data pre-processing and dimensionality-reduction strategies on clustering (Supp Notes 1.2). Two separate metrics were created: one to assess HLA haplotype-driven clustering (Global KNN_HLA_) and one to assess ancestry-driven clustering (Global KNN_ancestry_). *Global KNN_HLA_*: For a given individual, the KNN_HLA_ represents the average number of shared HLA alleles (out of 6) among the 10 nearest neighbors (maximum KNN_HLA_ = 6). The Global KNN_HLA_ is acquired by averaging KNN_HLA_ across the cohort. *Global KNN_ancestry_*: for a given individual, the KNN_ancestry_ represents the proportion of 10 nearest neighbors sharing the same ancestry label (maximum KNN_ancestry_ = 1.0). The Global KNN_ancestry_ is acquired by averaging KNN_ancestry_ across the cohort.

#### Quantification of CD8⁺ T cell escape across HLA haplotypes

Here, T-cell evasion across cohorts was analyzed using two distinct approaches: experimentally validated T cell evasion and predicted T cell evasion. For experimentally validated T cell evasion, we conducted an in-depth literature search to identify all studies that experimentally identified SARS-CoV-2 mutations leading to the reduction or abrogation of T cell response to epitopes. We identified 37 unique SARS-CoV-2 mutations leading to T cell evasion across 22 unique epitopes and 12 unique HLA alleles (Table S1). The resulting list of T cell evading mutations includes both HLA-evading as well as TCR-evading mutations. When available, we leveraged quantitative escape measurements, which we combined with published epitope immunogenicity (Response Frequency) scores to generate an immunogenicity-weighted escape score (Supp Notes 2.3). Since experimental studies are limited by the range of HLA alleles as well as SARS-CoV-2 variants investigated, we sought to complement this approach with a predictive framework. The latter leveraged the epitope-prediction tools netMHCpan-4.1^45^ (epitope-HLA prediction) as well as PRIME 2.1^46^ (epitope-TCR) to produce scores for both unmutated as well as mutated epitopes thereby predicting the impact of mutations on T cell immunity. Here, mutations causing percentile ranks to transition from strong HLA-binder (SB, %Rank < 0.5) to HLA non-binders (NB, %Rank > 2.0) were considered as leading to ‘epitope loss’. To reduce putative false positives commonly associated with predictive methods, only CD8+ T cell epitopes experimentally confirmed to be associated with an immune response (downloaded from IEDB, Jan 2^nd^, 2026) were retained for further analyses. To quantify total evasion per individual per HLA haplotype cluster, each HLA haplotype featured within the clustering analysis was associated with a total number of epitopes predicted to be abrogated by VoC mutations, considering all 6 classical alleles making up the HLA haplotype. In scenarios where participants possessed homozygous alleles, as well as alleles belonging to the same supertype, peptides were only counted once. As a result of this process, every participant was assigned a number corresponding to the total number of epitopes lost in the context of all variants analyzed (T-cell escape score). The average number of epitopes abrogated by VoCs across all HLA haplotypes of DBSCAN-defined clusters was determined, and inter-cluster variability was assessed using the Tukey test.

### Protein language model framework for mutation fitness estimation

#### Epistasis-aware Protein Language Model

We employed the model Evolutionary-Scale Modelling (ESM)-2^66^ , a transformer-based PLM trained on 65 million protein sequences acquired from the database UniRef^67^ . To obtain a *coronavirus*-specific model, we fine-tuned ESM-2 on sequences from *Coronaviridae* as previously described^51^. Briefly, we obtained the 650M parameters ESM-2 model (downloaded from https://github.com/facebookresearch/esm and manipulated through the Hugging Face Transformers library), and acquired *Coronaviridae* sequences from UniRef100 (Supplementary File 2). We leveraged a masked language learning strategy wherein 15% of randomly selected positions in each sequence were masked. Tokens (amino acids) were then predicted at each position in batched training (batch-size set to 5), with model weights updated by means of a cross-entropy loss function. The model was trained for 30 epochs with one epoch of warmup, as well as a base learning rate of 2e-5 controlled using a cosine-based learning scheduler. The success of fine-tuning was determined through perplexity scores, which correspond to the certainty with which the model predicts a masked token, with lower perplexity scores correspond to greater certainty of the prediction. Perplexity is defined as the exponential of the cross-entropy loss calculated during inference. Perplexity is defined by the following terms:

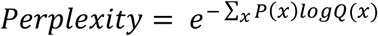

Where P(x) represents the true probability distribution and Q(x) represents the predicted probability distribution. Similar to values reported by Ito *et al* (2025), the out-of-the-box ESM-2 model possessed a perplexity of 11.38, whereas the model fine-tuned on *Coronaviridae* resulted in a perplexity of 1.17 when tested on coronavirus sequences. The resulting fine-tuned model was utilized to interrogate the fitness of mutations^50,51,53^ . Briefly, following tokenization of the query sequence, the position of interest was masked using a mask token. The fitness of any given mutation at that position can be computed by determining the probability of an amino acid occurring at that position given the surrounding context. In our case, we compared the log-likelihoods of the reference residue and the mutated residue at the relevant position^51,52^.

#### Fitness of viral mutations with ESM2-CoV

The resulting fine-tuned model was utilized to interrogate the fitness of mutations. Briefly, following tokenization of the query sequence, the position of interest is masked using a special mask token. The fitness of any given token (amino acid) at that position can be computed by determining the probability that a given sequence *x* would allow any token to occur at a given position *i* given the surrounding context and its corresponding constraints, θ (**Fig 5A**). Concretely,

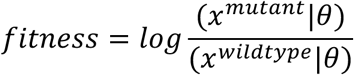

In our case, we compared the log-probabilities of the reference residue and the mutated residue at the relevant position.

#### Acquiring GISAID sequences for viral fitness and epistasis analyses

To interrogate the dynamic fitness of any mutation given an actively evolving virus, we fetched all viral sequences containing the mutation in question from GISAID and measured the fitness of that mutation against each observed mutational background. As a result, we obtained a dynamic fitness of a mutation as its surrounding sequence evolves. We downloaded all SARS-COV-2 Spike protein sequences from GISAID (as of October 30^th^ 2025, Supplementary File 3). All metadata was obtained through the GISAID interface using sample-specific EPI identifiers. Only sequences with a minimum length of 1100 amino acids and a minimum coverage of 95% were retained. To lower the computational burden, we made a subsampling of 50,000 sequences spanning the full dataset.

### Interaction maps between mutation pairs (query mutations and companion mutations)

ESM2-CoV was leveraged to generate mutation-pair interaction networks. Briefly, the fitness of each Omicron JN.1 mutation (query mutation) was estimated in the presence of a single other JN.1 mutation (companion mutation) for all JN.1 mutations to create a two-mutation interaction map. To achieve this, the Wuhan-1 spike protein sequence was first mutated with the companion mutation. The query mutation position was then masked, and the log-likelihood of the reference and mutated residues were acquired and used to compute the mutation fitness. Since the aim was to visualize the distribution of fitness effects of each query mutation across all companion mutations and not to compare query mutations to each other, all fitness metrics associated with one query mutation were normalized using a z-score ((data value – average of all points)/standard deviation).

## ACKNOWLEDGEMENTS

We acknowledge and thank GISAID as well as all contributing laboratories for giving access to their SARS-CoV-2 genome sequences. This study was completed thanks to computational resources provided by Compute Canada clusters Narval and Beluga. This study was also supported by funding from CHU Sainte-Justine, Canada Foundation for Innovation, IVADO COVID19 Rapid Response grant (CVD19-030), the Canadian Institutes of Health Research (CIHR) [#174924 and VR2-172712], the CIHR operating grant to the Coronavirus Variants Rapid Response Network [CoVaRR-Net, ARR-175622], and the COVID-19 Immunity Task Force/Public Health Agency of Canada. D.J.H. doctoral studies are supported by the Hydro Quebec Scholarship and Fonds de Recherche du Québec - Santé (FRQS) [350861]. J.G.H. holds a Tier 2 Canadian Research Chair from the NSERC [CRC-2024-00027].

## AUTHOR CONTRIBUTIONS

Conceptualization, D.J.H., E.C and J.G.H.; data curation and bioinformatic analysis, D.J.H., J-C.G.; Machine Learning, D.J.H. HLA-typing method development, data acquisition and processing, B.P., S.S., M.S.; contribution of RECOVER samples, H.D., B.B.; writing – original draft, D.J.H. and J.G.H; writing – review & editing, D.J.H., J-C.G., R.P.; B.B.; B.P.; S.S.; M.S.; H.D.; E.C.; J.G.H.; supervision, J.G.H., E.C., M.S., H.D.; funding acquisition, E.C., M.S., H.D. and J.G.H.

## DATA AVAILABILITY

This paper analyzes both existing, publicly available data as well as newly generated data. UKB is available upon request from https://www.ukbiobank.ac.uk/enable-your-research/apply-for-access. 1KGP HLA typing data is publicly available at http://ftp.1000genomes.ebi.ac.uk/vol1/ftp/data_collections/HLA_types/. HLA-typing data for the RECOVER cohort (generated here) are available in supplementary material (Supplementary File 2). All viral sequence data used in the fitness analyses were acquired from The Initiative for Sharing All Influenza Data (GISAID), at https://gisaid.org/ (Sequences found under EPI_SET_ID EPI_SET_260521gu). The user agreement for GISAID does not permit redistribution of sequences, but researchers can register to get access to the dataset. A GISAID acknowledgment table containing a full list of the laboratories and authors who contributed to the extensive GISAID SARS-CoV-2 genome database queried in this study is available in supplementary materials as (Supplementary File 3).

## CODE AVAILABILITY

All code used to perform analyses pertaining to HLA clustering, the comparison of HLA clustering methods, epitope loss predictions, as well as code used to fine-tune ESM-2, compute mutational fitness across sequence backgrounds, and visualize results are available on https://github.com/HussinLab/HLAScape. ESM-2 model weights and example usage (650M parameter esm2_t33_650M_UR50D model) can be found on https://github.com/facebookresearch/esm. NetMHCpan 4.1 and PRIME 2.0 command line tools and instructions for epitope predictions can be found on https://services.healthtech.dtu.dk/services/NetMHCpan-4.1 and https://github.com/GfellerLab/PRIME, respectively.

## EXTENDED DATA FIGURES

**Extended Data Fig 1.**
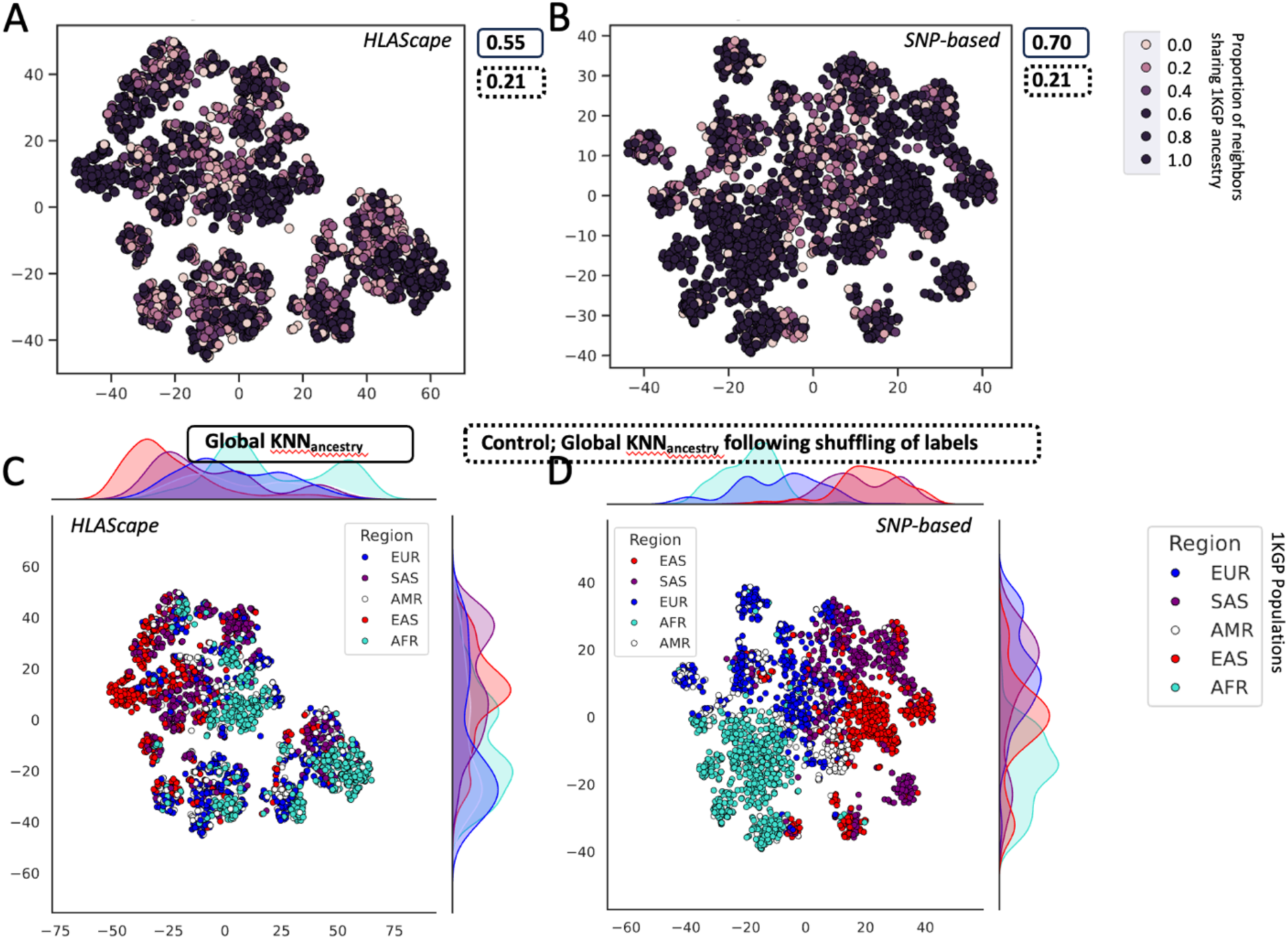
Impact of HLAScape and MHC SNP-based clustering of 1000 Genomes Project participants (n = 2693) on ancestry clustering. **A-B**. Clustering of 1KGP participants using HLAScape (**A**) or MHC-specific SNPs (**B**). Scatterplots are colored by Global KNN_ancestry_ (See Methods), with darker dots sharing ancestry label with a greater proportion of neighbors. **C-D**. Scatterplot coloring corresponds to 1KGP populations.

**Extended Fig 2.**
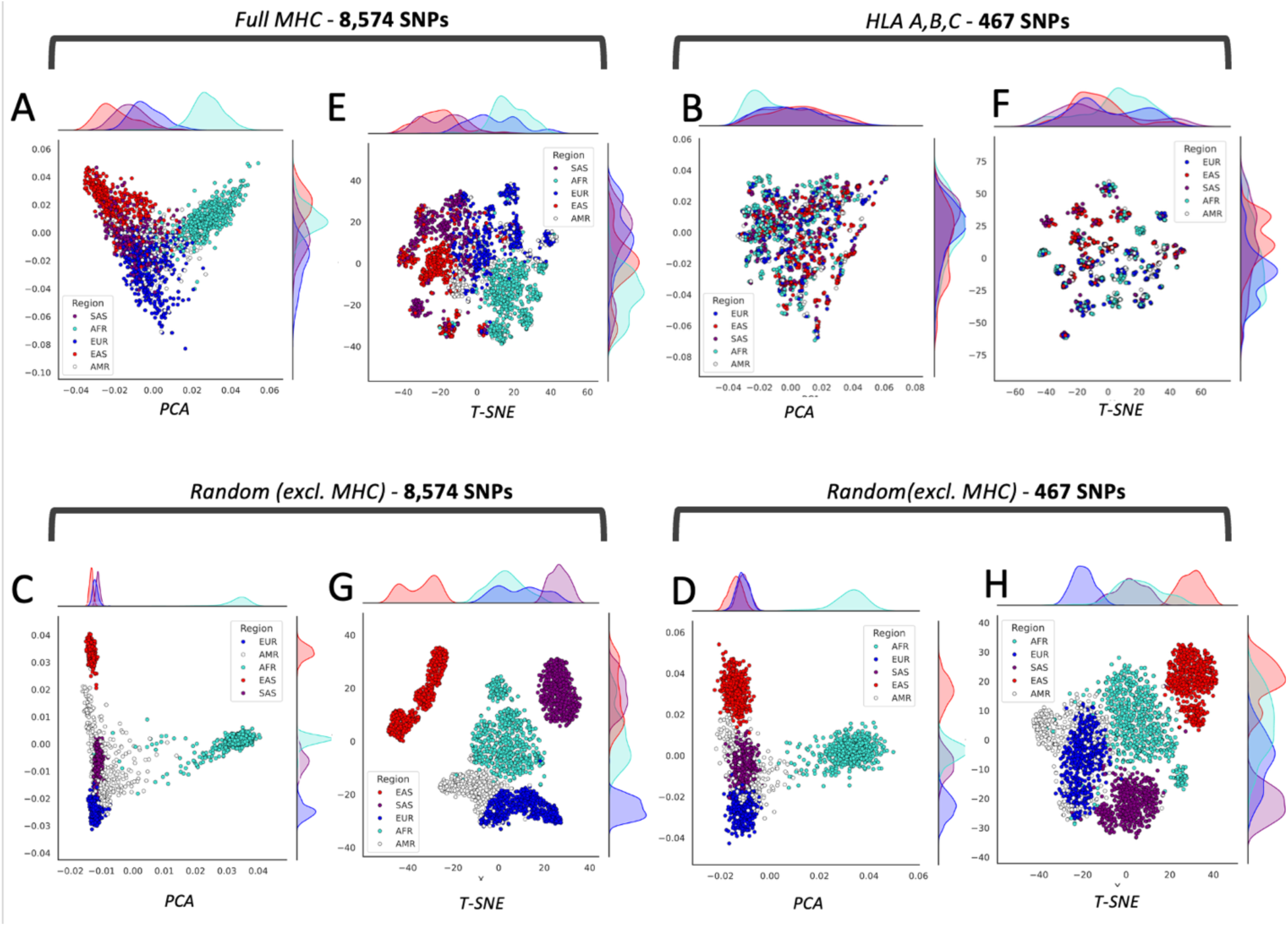
Summary of ethnicities across the 1000 Genomes Project dataset (n = 2693). **A-D**. PC1 v PC2 of SNPs identified within the MHC regions (MAF > 1%, LD R2 < 2 in blocks of 50 SNPs). SNPs were extracted from the complete MHC region, 8,574 SNPs (**A**); HLA A,B,C, 467 SNPs (**B**); Randomly-selected SNPs excl. MHC region (**C, D**). **E-H.** t-SNE on first 20 PCs of SNPs identified above.

**Extended Data Fig 3.**
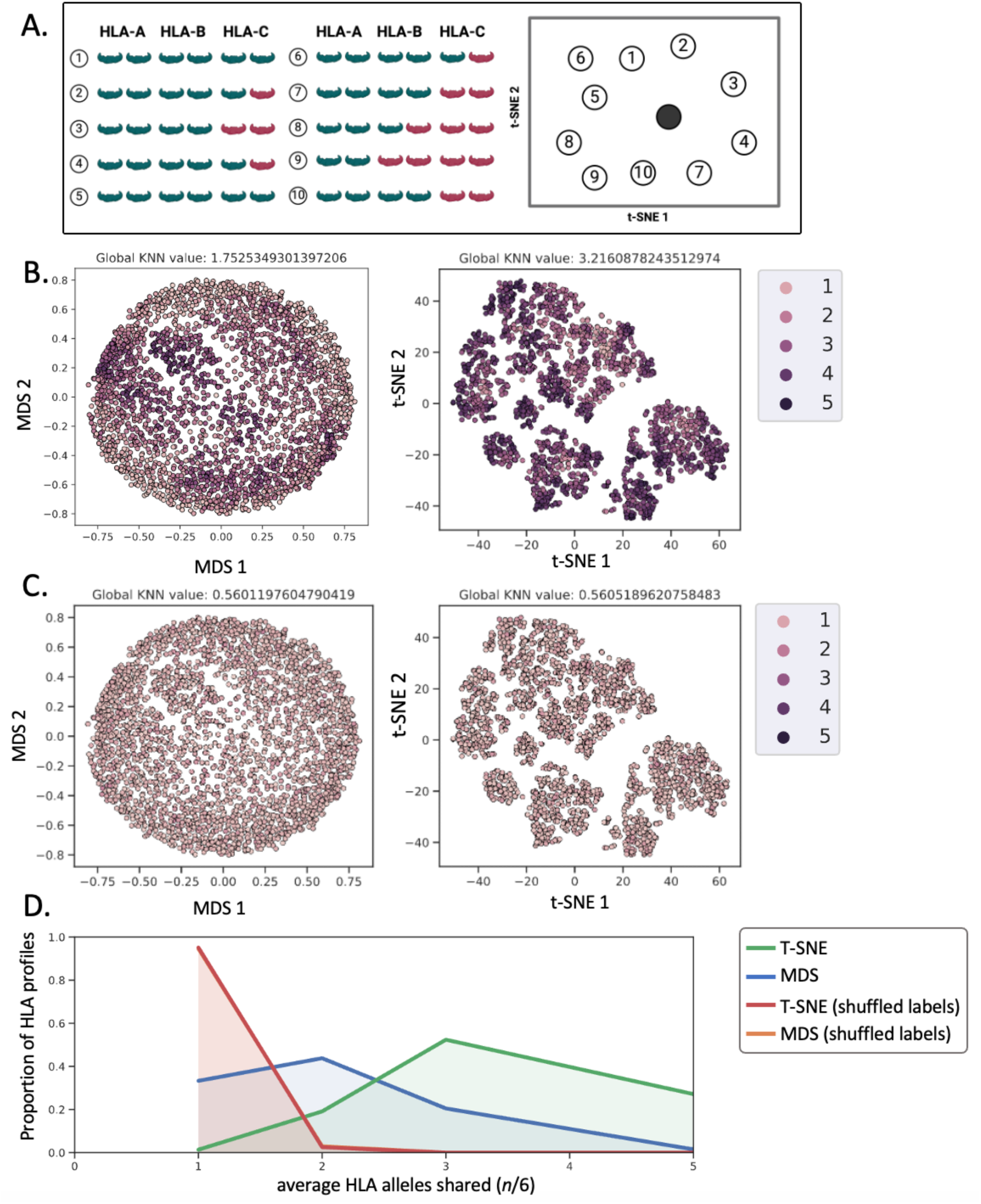
Validating HLA profile dimensionality reduction strategy through AI interpretability. **A.** K-nearest-neighbors (KNN) was utilized to establish an AI interpretability-driven metric to assess the degree of success with which different computational strategies clustered HLA profiles. This metric was used to validate the method utilized in HLAScape. Briefly, for a given clustered HLA profile, the average number of shared alleles across its 10 nearest neighbors was determined. Nearest neighbors were identified by computing Euclidean distance between dimensionality-reduced coordinates. **B.** The metric above was employed in the context of a subselection of UKB HLA profiles (n = 3000) clustered using t-SNE (left) or Multidimensional Scaling (MDS) (right). **C.** As a control, the analysis was repeated on the same dataset in which the labels were shuffled. **D.** Average number of HLA class I alleles (*n*/6) shared with 10 nearest neighbors across selected UKB HLA profiles (n = 3000).

**Extended Data Fig 4.**
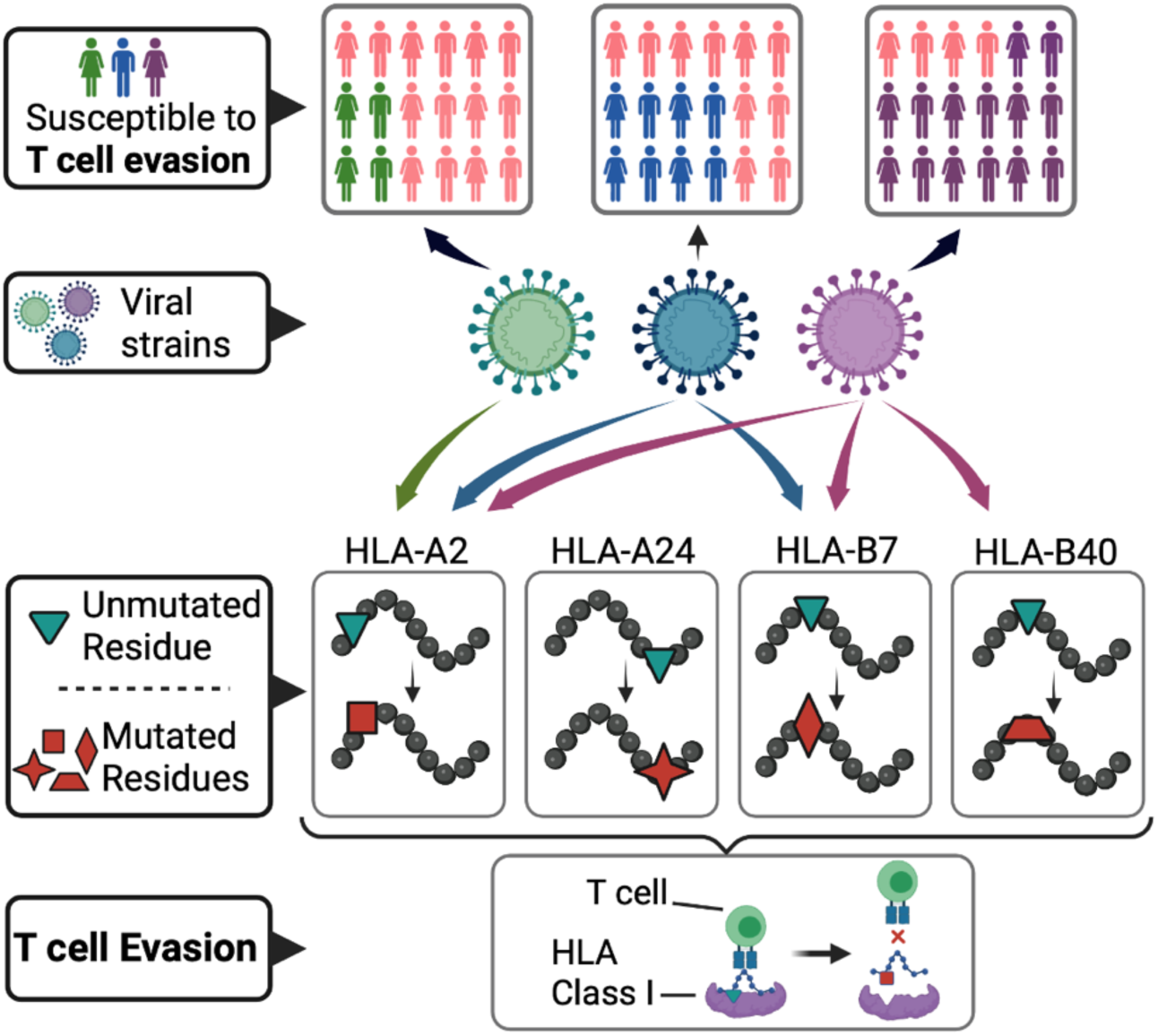
Identifying cumulative impact of variants on T cell evasion. We compile CD8+ T-cell epitopes with restricting HLA alleles and, for each viral variant, quantify predicted epitope disruption. We aggregate per-allele effects into a cumulative evasion score and project scores onto populations using HLA frequencies. This yields variant- and population-specific susceptibility profiles reflecting HLA diversity.

**Extended Fig 5.**
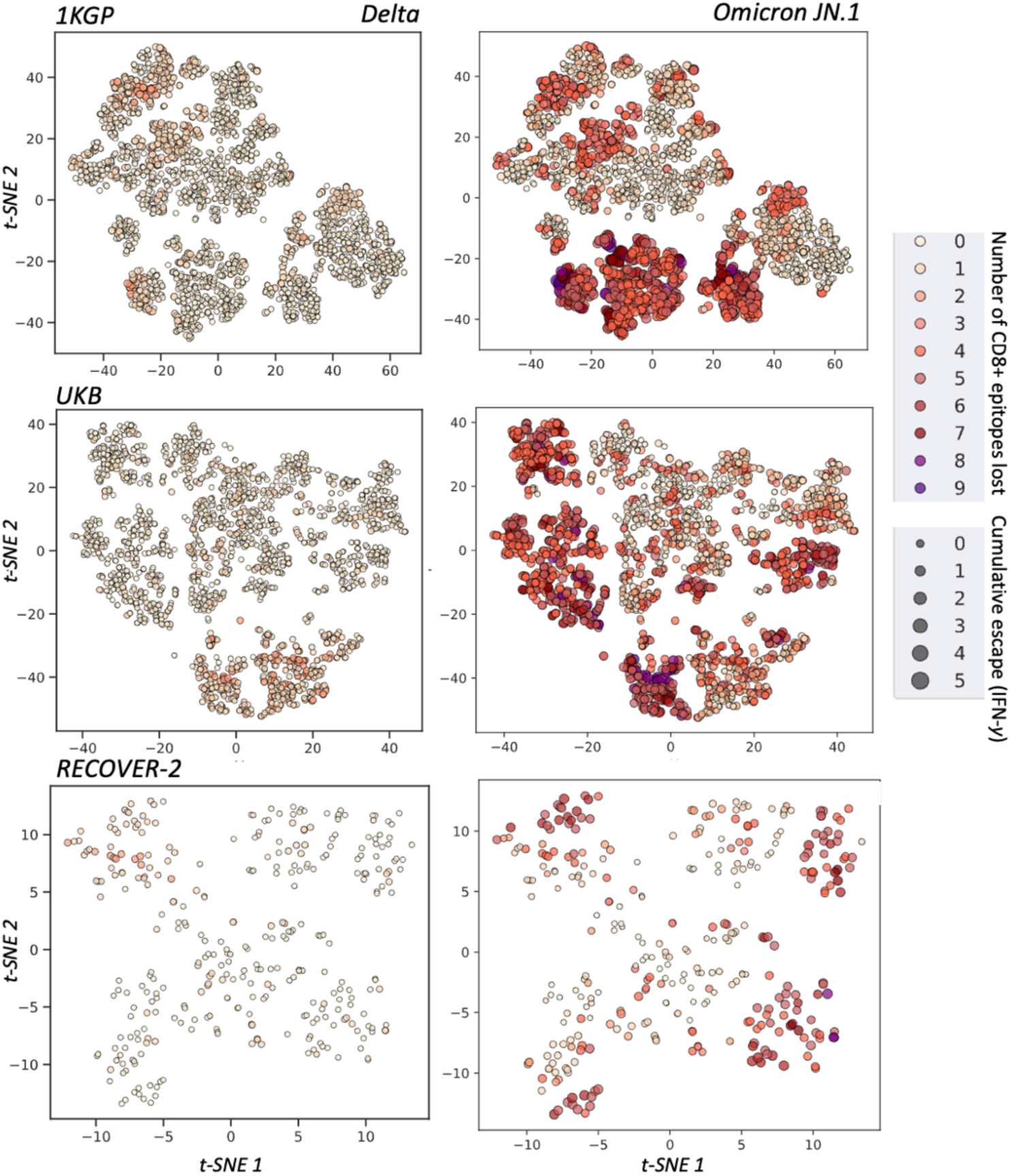
Experimentally validated and immunogenicity-weighted T cell escape across cohort HLA profiles. We compiled a list of peer-reviewed CD8+ T-cell epitopes escape mutations for quantitative escape values (reduction in measured IFN-*y*) were provided. We developed two scores to quantify escape: *(i)* number of epitopes lost *(shown as a color scale)*; and *(ii)* the overall impact on immune response as determined by multiplying the mutation-driven reduction in measured immune response (IFN-*y*) with the reported response frequency (proportion of patients with a positive response to the unmutated epitope), *(shown as dot sizes).* Results are shown for **(A)** 1KGP, **(B**) UKB, and **(C)** RECOVER-2. In each case, immunogenicity-weighted scores are shown in the context of mutations found in the SARS-CoV-2 VoCs Delta (left) and Omicron JN.1 (right)

**Extended Data Fig 6.**
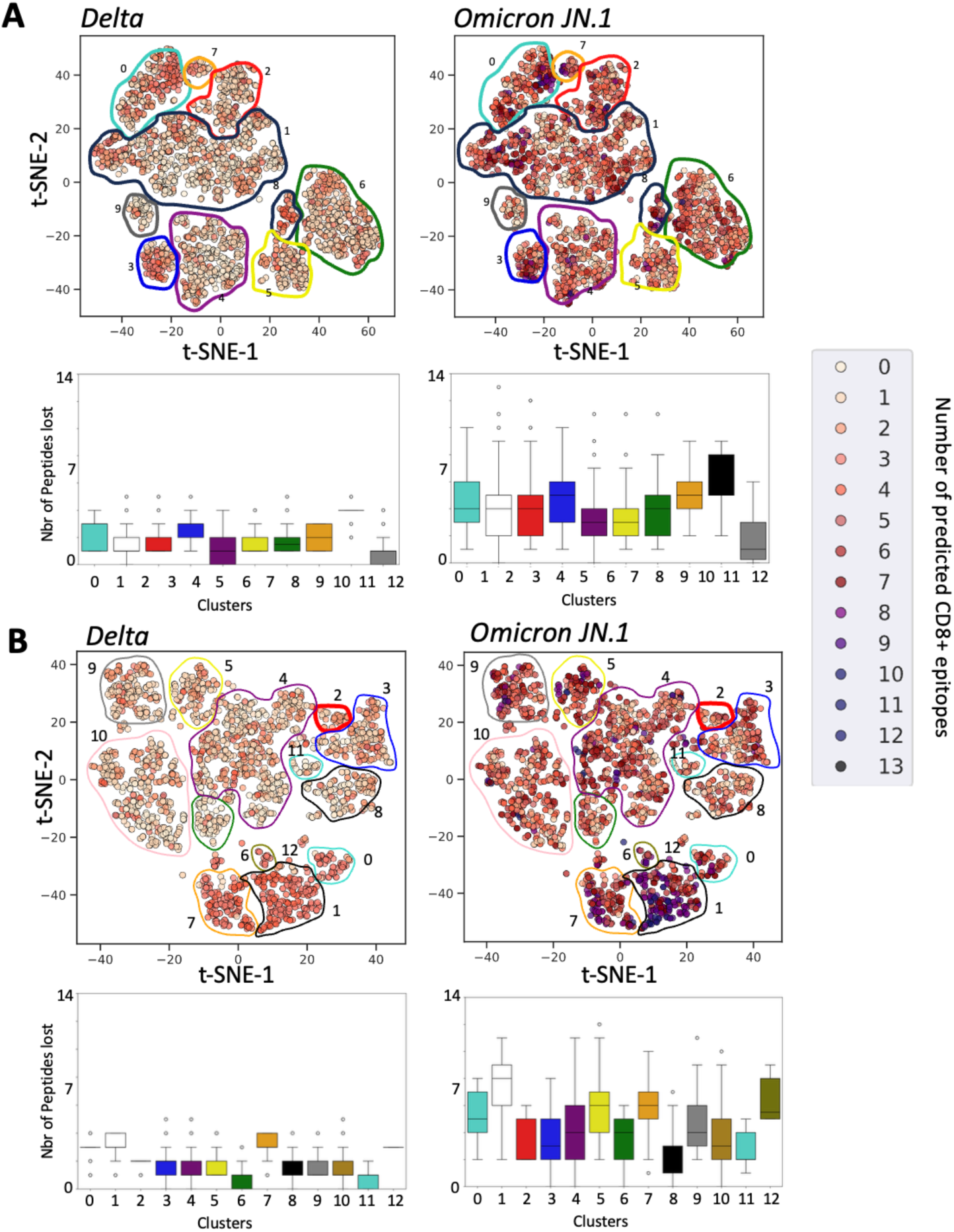
Predicted escape. Predicted immune evasion across cohort participants. Predictions were performed using netMHCpan 4.1 (peptide-MHC predictions) and PRIME 2.1 (peptide-MHC-TCR predictions). Predictions from both algorithms were combined to yield an evasion landscape reflecting both MHC evasion as well as TCR evasion (escape events predicted by both approaches were only counted once). Results are shown for 1KGP (**A**) as well as for the UKB (**B**).

**Extended Data Fig 7.**
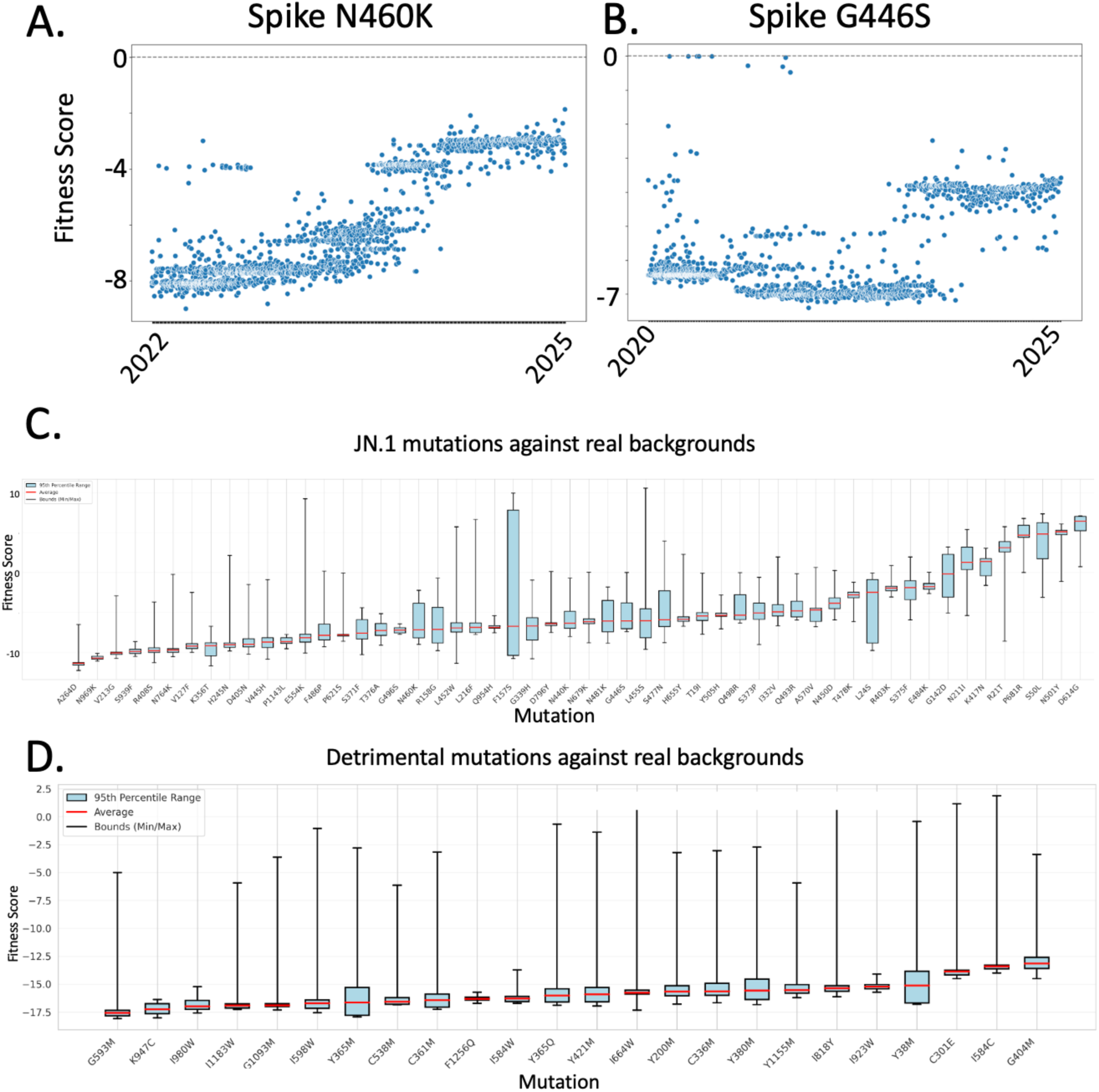
Tracking of viral fitness across pandemic. A, **B.** Estimated fitness of two illustrative mutations, Spike N460K **(A)** and Spike G446S **(B)**, against all backbones containing these mutations across a subsampling of 50K GISAID sequences spanning the pandemic. **C.** Summary representation of dynamic pandemic-wide fitness for JN.1 mutations. Here, each box represents the distribution of mutation-specific fitness across all relevant backbones (as in A. and B.) with the red lines, black lines, and blue boxes representing the average, distribution min/max, and distribution 95^th^ percentile range respectively. **D.** Estimation of fitness for low-fitness/low prevalence mutations against randomly-selected backbones spanning the pandemic.

**Extended Fig 8.**
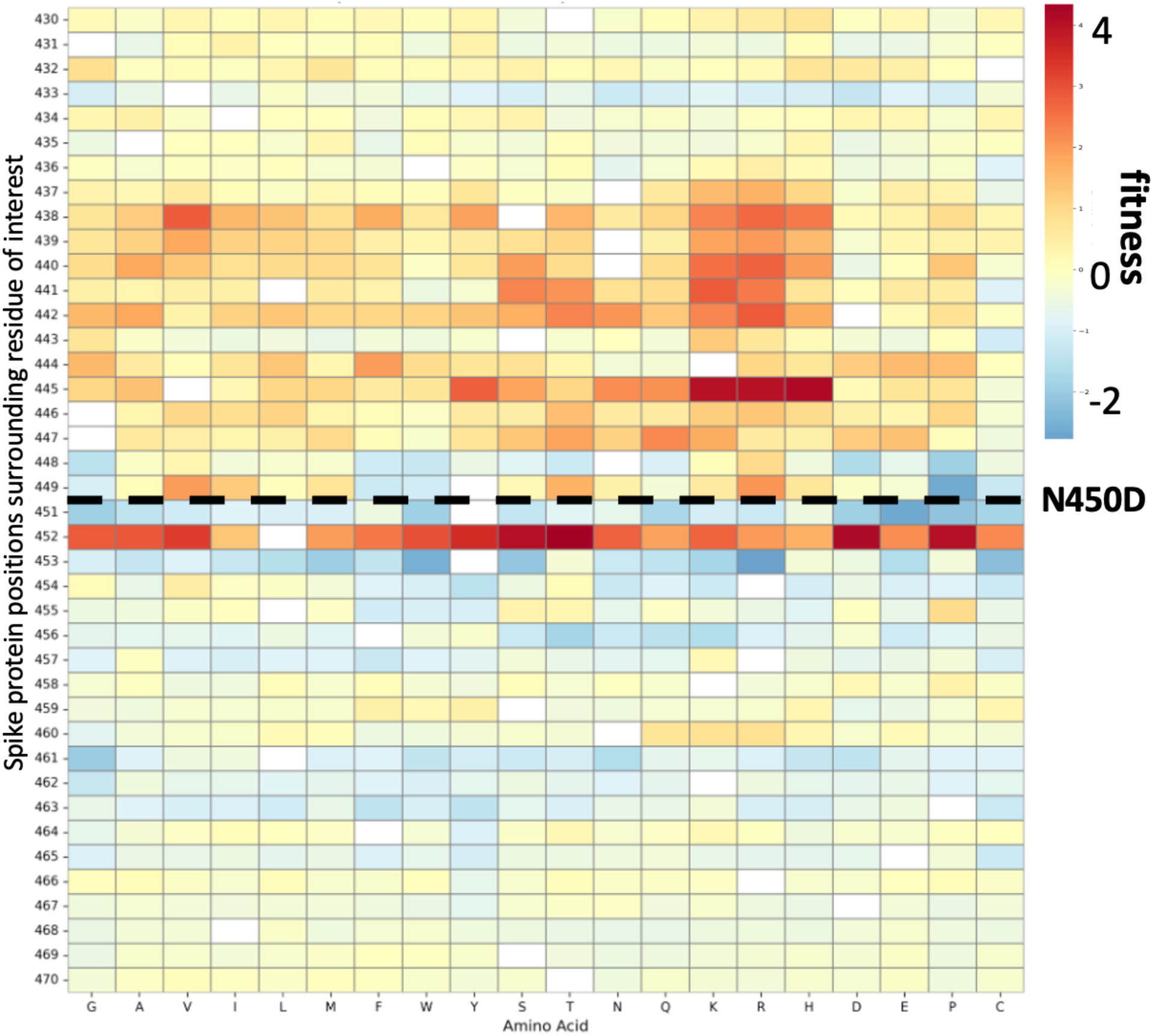
Interaction map between mutation of interest (Spike N450D) and all possible amino acid mutations across a -/+ 20 position window. The color bar corresponds to the fitness of the mutation of interest (Spike N450D) when in the presence of another mutation. All possible mutations were tested. Amino acids (x-axis) are sorted by chemical properties.

