## Supplemental_information for "Epistatic evolution drives HLA-dependent CD8+ T Cell escape risk in diverse populations"

### *1* Clustering strategy unveils discrepancies between ancestry- and HLA haplotype-generated groups

#### *1.1* Evaluation of HLA haplotype embedding and clustering

To evaluate clustering performance, we developed a K-nearest-neighbor (KNN)-based interpretability metric, which measures the average number of shared alleles across the 10 nearest neighbors for a given HLA profile (**Ext. Data Fig 3A**). Results pertaining to the 1KGP are shown in **Fig 1C**. Here, the Global $KNN_{HLA}$ corresponds to the average number of HLA alleles (over a total of 6 alleles making up classical class I haplotypes) shared across all 10 nearest neighbors, averaged across the cohort. Individuals across the 1KGP share on average 3.21 HLA alleles with their nearest neighbors according to our clustering scheme. As a negative control, labels were shuffled, resulting in little to no clustering based on HLA profile similarity. This is confirmed by the control Global KNN, where individuals shared only 0.57 HLA alleles with their nearest neighbors.

#### *1.2* Comparison of dimensionality reduction and preprocessing

##### *1.2.1 Comparison of HLAScape with classical population genetics approach.*

We compared the HLA mapping strategy proposed here to a classical SNP-based approach (**Fig 1C-E, Ext. Data Fig 2, Fig S1**). Briefly, SNPs were extracted from HLA alleles A, B and C across the 1KGP (see methods), and clustered by performing t-SNE on the first 20 Principal Components generated from the SNP genotypes. This approach resulted in a slightly lower Global $KNN_{HLA}$ value (2.76) despite a comparable control Global $KNN_{HLA}$ (0.58), suggesting that SNPs extracted the HLA region may not be as representative of populational HLA structure as phased HLA types. To ensure that the number of SNPs extracted was sufficient, this analysis was repeated by randomly selecting a similar number of SNPs across chr6 (excluding the MHC region) (**Fig 1E**). As expected, the exclusion of the MHC region resulted in a negligible Global $KNN_{HLA}$ of 0.92.

It is interesting to note that broader windows within the MHC regions (ex. Full set of SNPs across the MHC locus) resulted in a stronger populational signal (**Fig S1**). This method resulted in a populational structure similar to that reported by others (**Fig S1**) [1] , validating our SNP-based analysis. This signal was partially degraded in more targeted peptide presentation genes (HLA class I A, B, C), although clustering by HLA labels improved as the SNP-extraction window targets such genes. Nevertheless, the strongest clustering occurs when directly leveraging functional HLA labels. In contrast, applying 1KGP ancestry labels to the HLAScape-grouped individuals showed modest clustering of populations, with East Asian (EAS) and African (AFR) populations demonstrating the strongest clustering.

##### *1.2.2 Comparison of Jaccard Distance + t-SNE vs PCA + t-SNE vs t-SNE alone*

To determine if the clusters and clustering efficiencies achieved were method-specific (t-SNE), we compared the clustering approach featured in HLAScape (Jaccard distance + t-SNE) to alternative strategies (**Fig S2**). We explored alternative t-SNE pre-processing strategies, which included Principal Component Analysis (PCA) as well as t-SNE applied directly to the binarized HLA dataset. While all methods resulted in significant grouping of similar HLA profiles, the Jaccard distance + t-SNE based approach yielded superior clustering, with a Global $KNN_{HLA}$ value of 3.21. The use of t-SNE on PCA-reduced data, as well as t-SNE directly applied to binarized HLA data, yielded inferior Global $KNN_{HLA}$ values of 3.08 and 0.75, respectively.

##### *1.2.3 Comparison between t-SNE vs DMS*

To assess the relevance of t-SNE, we explored Multidimensional Scaling (MDS), an alternative dimensionality reduction method. Again, MDS yielded an inferior Global $KNN_{HLA}$ value of 1.75 (**Ext. Data Fig 3B**). Nevertheless, further interrogating the HLA composition of the clusters obtained using MDS resulted in clusters highly similar to those obtained by t-SNE, suggesting the reproducibility of results across both methods. In all approaches tested, the negative control (shuffled HLA alleles) resulted in $KNN_{HLA}$ values much lower than their experimental counterparts, suggesting a consistent ability to group individuals by HLA profiles. This resulted in a significantly lower $KNN_{HLA}$ values below ($< 1$) (**Ext. Data Fig 3B, C**). Collectively these results indicate that the clusters obtained are not method-specific, although the grouping of similar HLA haplotypes is more efficient when relying on Jaccard Distance + t-SNE.

#### 2 Population-scale HLA-typing and visualization of a real-application cohort, RECOVER cohort (n = 576).

##### 2.1 HLA-typing and visualization of the RECOVER cohort

To investigate T-cell evasion across an HLA-diverse population, our computational strategy was applied to the RECOVER cohort, a cohort of SARS-CoV-2 convalescent individuals. To enable HLA profile mapping with HLAScape, the cohort was entirely HLA typed utilizing a high-throughput, cost-effective nanopore long-read sequencing workflow (**Fig 3A**). Peripheral blood mononuclear cells (PBMCs) were collected from RECOVER participants (**Fig 3A**, left), and targeted long-read sequencing of HLA class I loci was performed by adapting the strategy described by Stockton et al, 2020 [2] for 96-well plate processing and laboratory automation (**Fig 3A**, center). This adaptation enabled parallel processing of 96 samples per batch, including automated reaction setup, post-PCR normalization, and pooling. Amplicons were quantified in plate format using a fluorescence-based dsDNA assay, and concentration values were used to normalize and pool samples using an automated liquid-handling workflow prior to library preparation. Across sequencing batches, libraries were prepared using Oxford Nanopore ligation-based (SQK-LSK109 with EXP-NBD196) and rapid barcoding (SQK-RBK111.96) workflows. Sequencing was performed on MinION R9.4.1 flow cells, and HLA class I genotypes were inferred using HLA*LA software (Dilthey et al., 2019) [3] (**Fig 3A**, right). Only patients for which all 6 HLA class I alleles were identified with high confidence (per-allele quality score, Q1, above 0.8) were retained for further analyses (n = 334, Supplementary File 1).

This combined experimental and computational workflow enabled scalable HLA typing in a clinical cohort and supported downstream analyses of HLA haplotype diversity and T-cell escape. Before performing escape analyses, we confirmed that the HLA profile distributions of the UKB and RECOVER cohorts were highly similar, as shown by their co-clustering and comparable allele-frequency patterns (see Fig 2B and 2C). Application of the HLAScape workflow to the RECOVER cohort identified 7 distinct HLA haplotype clusters, numbered 0 to 6 (**Fig 3D**).

##### 2.2 Comparison of HLA haplotype maps across cohorts

Prior to conducting analyses pertaining to T-cell evasion, we sought to establish a comparative analysis between the HLA profiles specific to the UKB and those identified within the context of the RECOVER-2 cohort. The HLA profile clustering feature of HLAScape was applied to a collection of HLA profiles comprising individuals from both the UKB and the RECOVER-2 cohorts (**Fig 3B**). All RECOVER-2 HLA profiles co-clustered with UKB HLA-profiles (purple and white dots, respectively), resulting in no clusters specific to either cohort, suggesting a high level of similarity between the HLA profiles of both cohorts. This similarity is further supported by the comparison of HLA allele-specific frequencies across both populations (**Fig 3C**), thus indicating significant overlap between the HLA allele distributions of both cohorts and establishing the comparability of the two cohorts.

##### *2.3* Leveraging published quantitative escape data for an immunogenicity-weighted score

We curated experimentally supported human CD8+ T-cell epitopes with reported restricting HLA class I alleles and, for each SARS-CoV-2 variant, annotated epitope-overlapping mutations using studies that provided quantitative mutant vs wild-type (WT) functional readouts. For each epitope–mutation pair, we defined a residual response ratio (*r*) as mutant/WT response. For dose–response datasets, *r* was computed as the ratio of areas under the response curves across the tested peptide concentration range ($AUC_{Mut}$/ $AUC_{WT}$); for donor-level datasets without dose titrations, *r* was computed as the mutant/WT ratio at matched conditions (aggregated as reported). We then quantified variant-specific susceptibility per HLA genotype using two complementary metrics: (i) the number of epitopes with reported reduced recognition (visualized as a color scale), and (ii) an immunogenicity-weighted impact score $I = (1 - r)\ x\ f$ where (f) is the epitope response frequency from IEDB (proportion of tested subjects responding to the unmutated epitope), visualized as dot sizes. This framework integrates mutation-driven functional escape with population-level epitope immunogenicity to generate variant- and population-specific profiles reflecting HLA diversity.

##### *2.4* Generation of Synthetic SARS-CoV-2 variants

200 synthetic SARS-CoV-2 variants were generated *in silico* to retain the evolutionary mutational structure observed in Omicron sublineages. For each synthetic variant, novel mutated positions were randomly selected proximally to existing Omicron BA.2 mutations (+/- 20 amino acid positions) and were mutated to randomly selected amino acids. To generate synthetic variants that completely lacked the mutational structure observed in Omicron lineages, 200 additional synthetic variants were generated by randomly selecting 200 positions across the SARS-CoV-2 proteome and mutating them to a randomly selected amino acid. All resulting synthetic SARS-CoV-2

variants were analyzed using HLAScape in the context of HLA profiles obtained from the UKB biobank and the RECOVER-2 cohort.

#### *3* Leveraging an epistasis-aware Protein Language Model to forecast T-cell evading mutations

##### *3.1* Validating single-mutation and epistatic fitness predictions of the fine-tuned model

As an initial gage of the ability of our fine-tuned model's ability to estimate the fitness of single mutations, we computed the fitness of all possible point mutations across the Spike protein and compared these values to epidemiological data. Briefly, we subsampled pandemic-wide SARS-CoV-2 sequences (n = 49,921, see Methods for data pre-processing, Supplementary File 3), and computed the number of occurrences of each mutation across this subsampling, and the resulting frequency with the ESM-2-estimated fitness (**Fig S8A**). We obtained a positive relationship between these two values, suggesting that ESM-2 can successfully identify beneficial viral mutations (**Fig S8A**).

To assess epistasis-dependant fitness, we analyzed DMS data from Bloom *et al* . In this study, Bloom *et al* [4] conducted a DMS experiment on the SARS-CoV-2 Spike protein, using variants BA.2 and XBB 1.5 as mutational backbones, yielding an ideal dataset to validate our method. We used our fine-tuned model to estimate the fitness of all mutations featured within the datasets (every possible Spike missense mutation) either in the context of the Wuhan-1 Spike sequence or the assayed mutated Spike sequence (BA.2 or XBB). Mutations that scored low in DMS experiments were estimated to have low fitness when predicted in the context of either the Wuhan-1 or BA.2/XBB 1.5 sequences (**Fig S8B-D**). In contrast, mutations that scored higher in the DMS experiments were estimated to have higher fitness when assessed against a variant backbone compared to a Wuhan-1 backbone, suggesting the ability of our fine-tuned model to capture mutational interactions (**Fig S8B-D**).

##### *3.2 Background-dependent fitness of SARS-CoV-2 mutations across the pandemic*

Here, we sought to investigate the ability of our framework to dynamically assess the fitness of viral mutations throughout the pandemic across many viral sequences. To do this, we utilized sequences from the viral database GISAID (Supplementary File 3). For each mutation of interest, we collected all high-quality (>95% coverage) sequences containing that mutation. We then estimated the fitness of the mutation against all collected background. Initial focus was given to known highly successful viral mutations. Our analysis of the Spike mutations N460K and G446S show gradual increase in fitness as the surrounding sequence evolves, showcasing a highly dynamic background-dependant fitness (**Ext. Data Fig 7A-B**). We expanded our analyses to all mutations associated with select variants, namely Delta, Omicron BA.2, and Omicron JN.1. For each variant, the fitness of each mutation was computed against all observed backgrounds, and averaged (**Ext. Data Fig 7C**). As expected, none of the mutations featured here are detrimental when averaged across sequences, although average fitness estimates range from neutral to beneficial. To compliment this, we sought to determine whether the model overly relied on beneficial sequence backgrounds, regardless of the mutation of interest. Here, we hypothesized that mutations pre-determined to have low fitness will be determined as such even when placed in a beneficial background To test this, we selected a set of mutations that we defined as having low fitness using the following criteria: they were never observed in the pandemic, and they scored low on all functional DMS assays. These were inserted in prevalent variant (Variants of Concerns, VoC) backgrounds, and their fitness was estimated. All mutations remained highly detrimental despite a beneficial mutational background (**Ext. Data Fig 7D**).

##### *3.3 Interaction maps between triple (query mutations and companion mutations pairs)*

ESM2-CoV was leveraged to generate mutation-triplet interaction networks. The fitness of each Omicron JN.1 mutation was estimated in the presence of two other JN.1 mutation for possible JN.1 3-mutation combinations to create a three-mutation interaction map. To achieve this, the Wuhan-1 spike protein sequence was first mutated with the two companion mutations. The query mutation position was then masked, and the log-likelihood of the reference and mutated residues were acquired and used to compute the mutation fitness (see Methods). We ensured that the query mutation never corresponded to one of the companion mutations. Since the aim was to visualize the distribution of fitness effects of each query mutation across all companion mutations (and not to compare query mutations to each other), all fitness metrics associated with one query mutation was normalized using a z-score ((data value – average of all points)/standard deviation). Results were summarized as an interaction heatmap (**Fig S10**) in which the y-axis represents each query mutations and the x-axis represent the companion mutation pair. Using this approach, we observe a wide array of varying inferred epistatic interactions ranging from both proximal and distal interactions. These results allow us to disentangle epistatic interactions. For example, the mutation

K417N has high fitness when in the presence of mutations N440K and N460K but has lower fitness in the presence of either mutations individually or combined with different third partners (**Fig S10**).

##### *3.4 Exploring the putative mutational space within epitope-specific mutational hotspots*

Two spike mutational hotspots previously implicated in CD8+ T-cell escape were selected for focused analyses: (i) the JN.1 hotspot comprising N450D, L452W, and L455S, and (ii) the BA.2/JN.1 hotspot comprising S371F, S373P, S375F, and T376A. Each hotspot was linked to an experimentally characterized loss of recognition for an immunodominant HLA-A*24:02–restricted epitope (NYNYLYRLF / NYDYWYRSF for the N450/L452/L455 region; and LYNSASFSTF / LYNFAPFFAF for the S371–T376 region). These epitope–HLA assignments and their associated escape evidence were taken from the experimental source(s) used in this study and recorded for downstream interpretation. To explore a broader space of potential epistatic partners beyond observed combinations, an exhaustive local scan was performed for each hotspot mutation. Briefly, for each hotspot mutation, all possible single–amino acid point mutations were introduced one-at-a-time across a 40-position window centered on the hotspot region (e.g., ±20 residues around the hotspot; window boundaries defined relative to spike indexing). At each position in the window, all non-reference amino acid substitutions were considered (typically 19 alternatives per site), yielding a comprehensive library of "query mutation + companion mutation" double mutants. Each double mutant was scored using the same fitness framework described above, and interaction scores were computed relative to the corresponding single mutants and background. Results were summarized as local interaction maps, highlighting combinations that were predicted to be high-fitness and/or strongly epistatic (**Ext. Data Fig 8**, **S11–S12**). Alongside recapitulating the beneficial effect of known hotspot mutations on the mutations of interest, we identify additional non-observed high-fitness mutation pairs, thus suggesting potential routes for further expansion of these hotspots.

#### *4* Supplementary figures

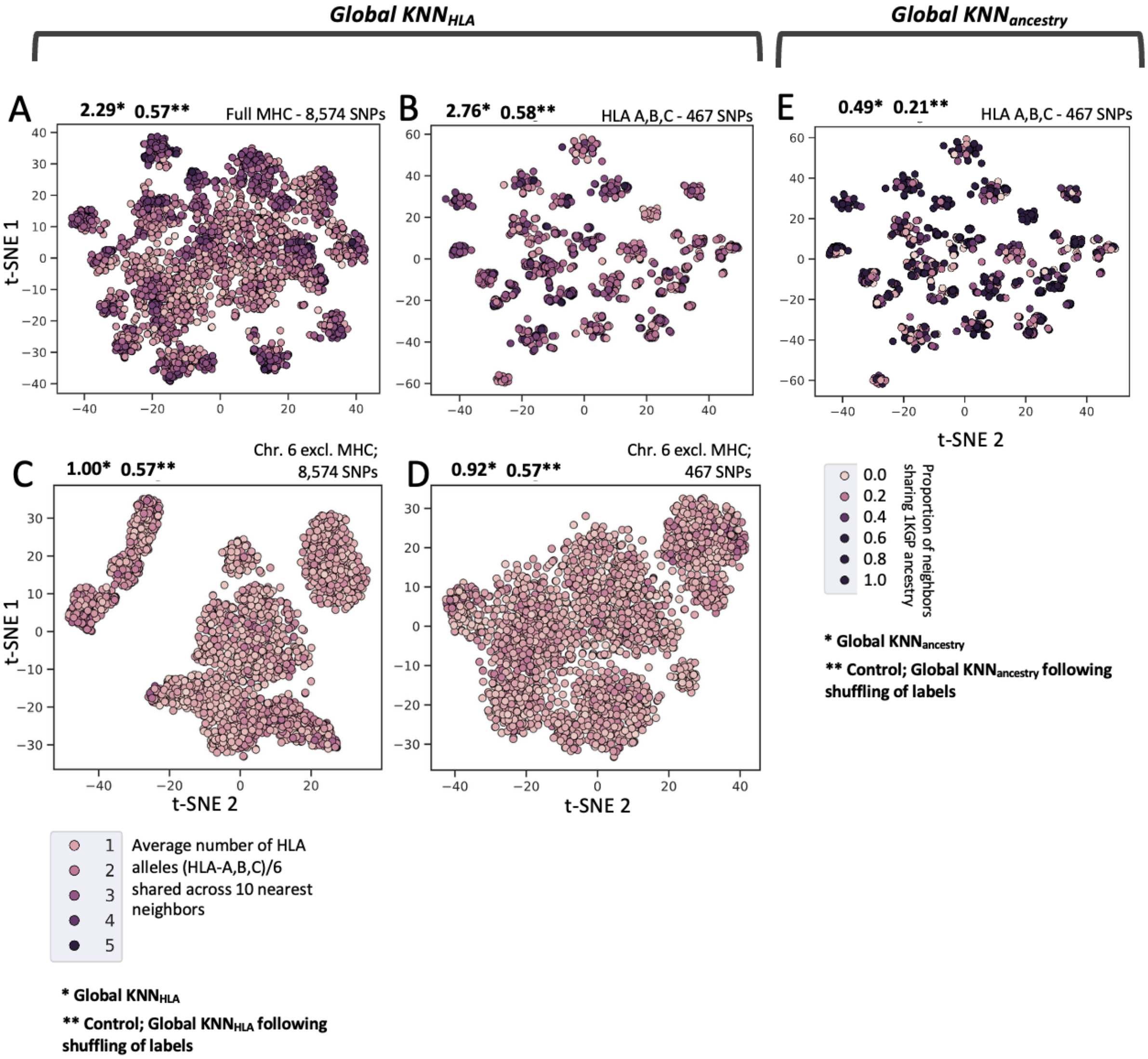

**Fig S1.** KNN-based interpretability metric for HLA haplotype clustering, measuring the average number of shared alleles across the 10 nearest neighbors for a given HLA profile. Global $KNN_{HLA}$ (*) is the average number of shared HLA alleles (out of 6) among the 10 nearest neighbors, across the cohort; control (**) is the Global $KNN_{HLA}$ following shuffling of labels. **(A)** SNPs were extracted from the complete MHC region, 8,574 SNPs; **(B)** HLA A,B,C, 467 SNPs; **(C)** Randomly-selected SNPs (excl. MHC region) matching SNP count for full MHC region (8,574 SNPs); **(D)** Randomly-selected SNPs (excl. MHC region) matching HLA A,B,C (467 SNPs). In all cases, SNPs were clustered using t-SNE on the first 20 Principal Components from SNP genotypes. (**E**). Global $KNN_{ancestry}$ metric applied to clustering of 1KGP based on HLA A,B,C (467 SNPs).

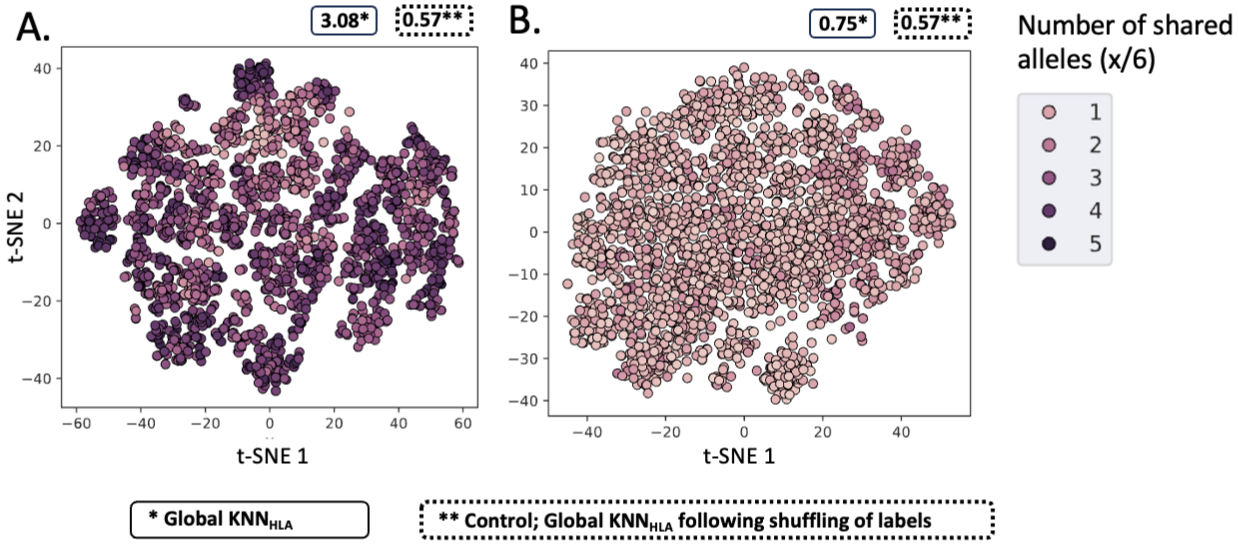

**Fig S2. Validating the HLA profile pre-processing strategy for clustering through AI interpretability. A.** HLA-map of 1KGP following HLA profile preprocessing by PCA (20 components) instead of Jaccard distances prior to t-SNE. **B.** HLA-map of 1KGP with t-SNE applied directly to binarized HLA profiles

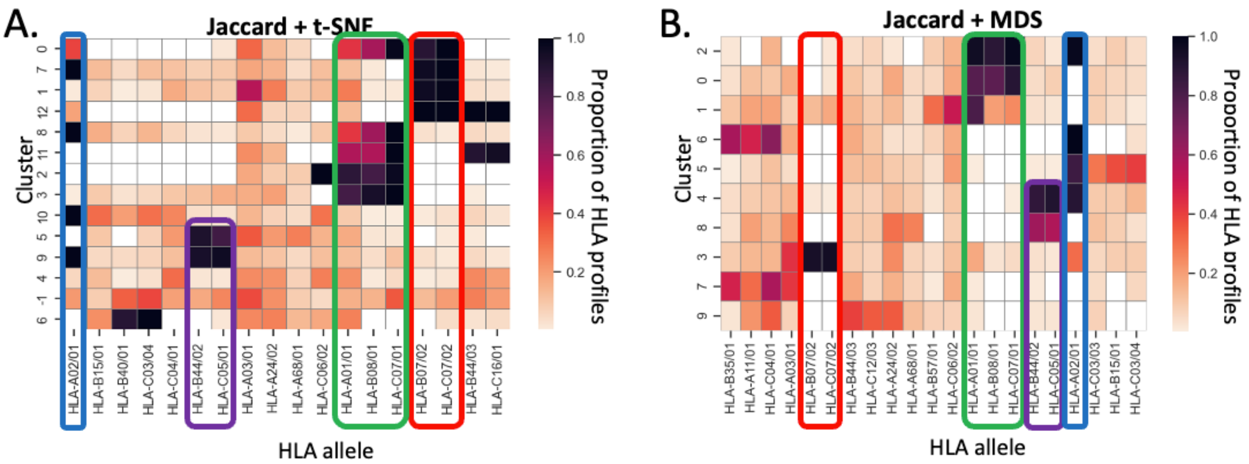

**Figure S3. Validating the HLA profile clustering strategy through AI interpretability. A, B**. Top four HLA alleles are shown for each cluster for Jaccard + t-SNE as well as Jaccard + MDS. Clusters (y-axis) were identified by DBSCAN. Both clusters and HLA alleles were ordered using hierarchical clustering to reflect the clustering structure achieved by the dimensionality reduction technique in question (t-SNE, MDS).

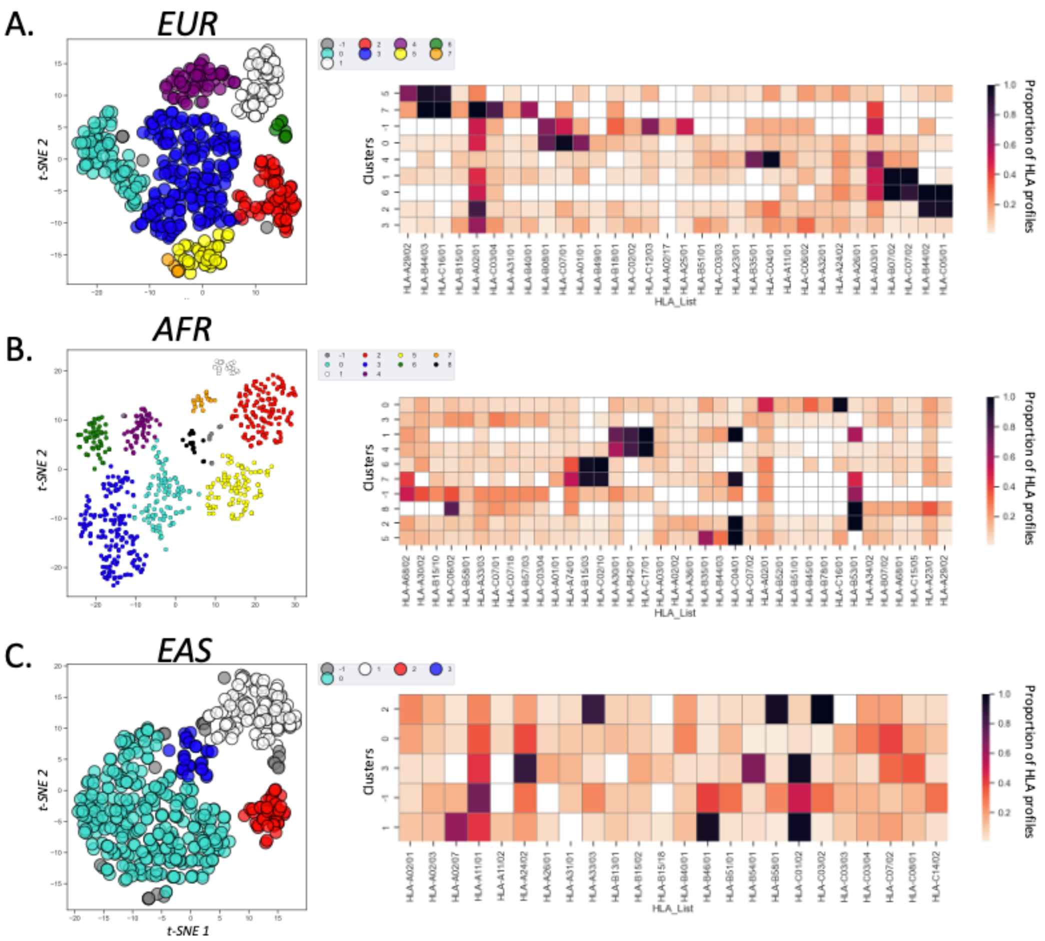

**Figure S4. predicted epitope loss across ethnicities.** Here, predicted epitope loss across HLA profile-defined clusters are shown across European (EUR, top), African (AFR, middle) and East Asian (EAS, bottom) populations, acquired from the 1000 Genomes Project.

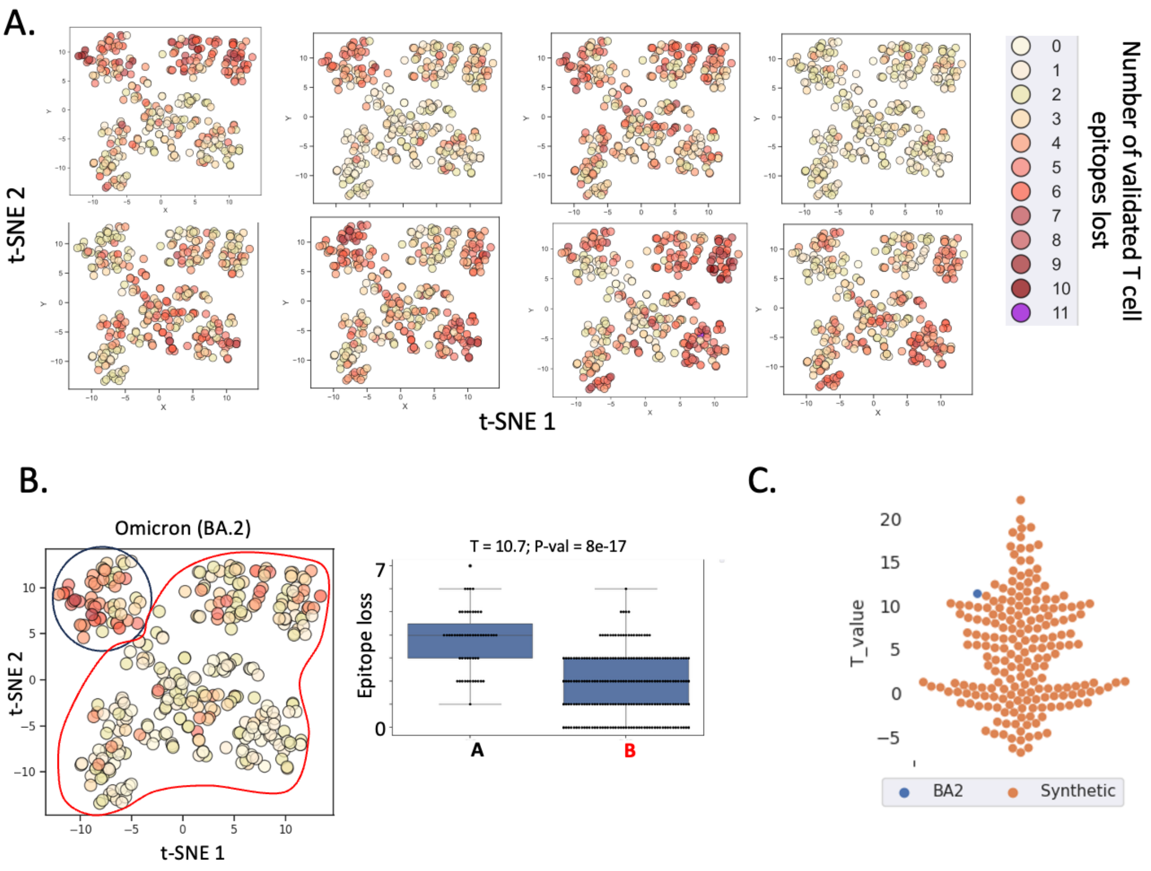

**Figure S5. Epitope loss by synthetic variants in the RECOVER cohort**. **A.** Impact of synthetic mutants on epitope loss across the RECOVER cohort**.** 200 variants were synthesized to mimic the mutational hot spots of VoC Omicron BA.2 (8 synthetic variants shown here). Briefly, synthetic mutants were generated by creating novel mutations at near the original BA.2 mutations. Both the distance from the original mutations (ranging from -20 to +20 amino acids) as well as the mutated residue were randomized. **B.** Example of T test comparing predicted T cell evasion by SARS-COV-2 mutations (example given for Omicron mutations BA.2) in the context of RECOVER HLA cluster #1 and the remainder of the cohort. **C.** distribution across synthetic variants (orange dots) of T values obtained from t-tests comparing epitope loss in cluster #1 to other clusters. These are compared to the t value obtained from BA.2 (blue dot).

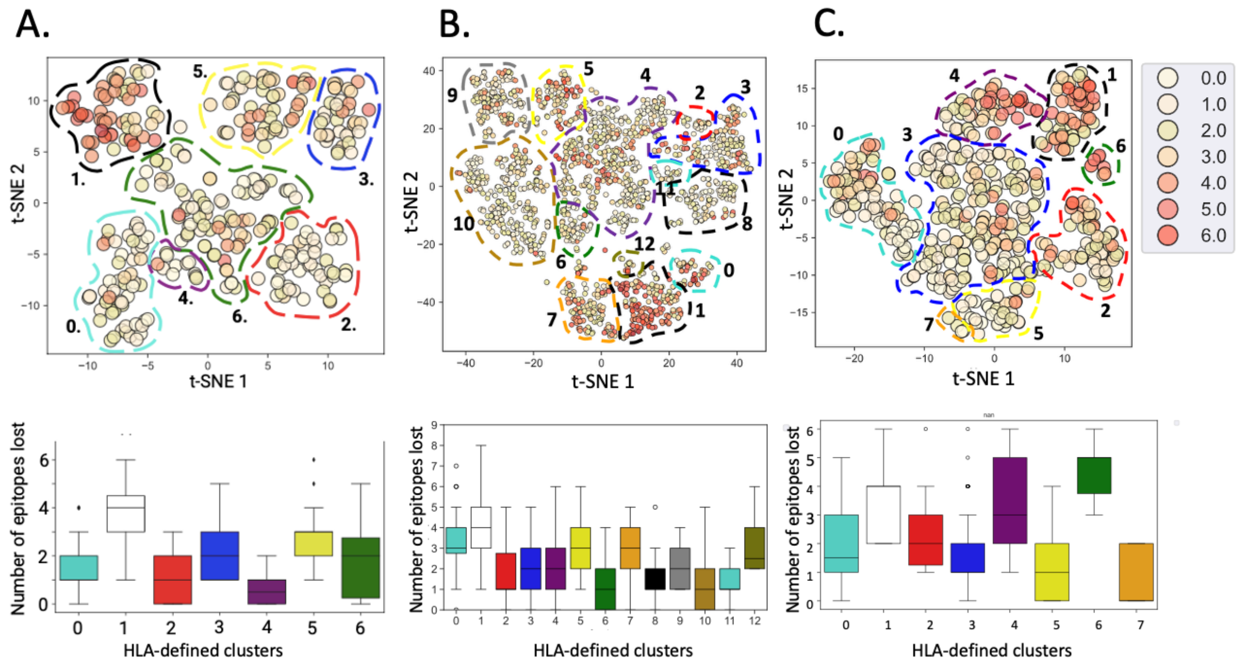

**Figure S6. CD8+ T cell epitope loss across cohorts. A-C.** Predicted CD8+ T cell epitope loss across the RECOVER cohort; the UKB, and the 1KGP European population. Dots (center figure) correspond to unique HLA profiles (HLA-A, B and C), clustered by HLA-allele similarity. The color scheme corresponds to the number of epitopes predicted to be lost by Omicron (BA.2) mutations. Boxplots correspond to the average number of CD8+ T cell epitopes lost per HLA-defined cluster.

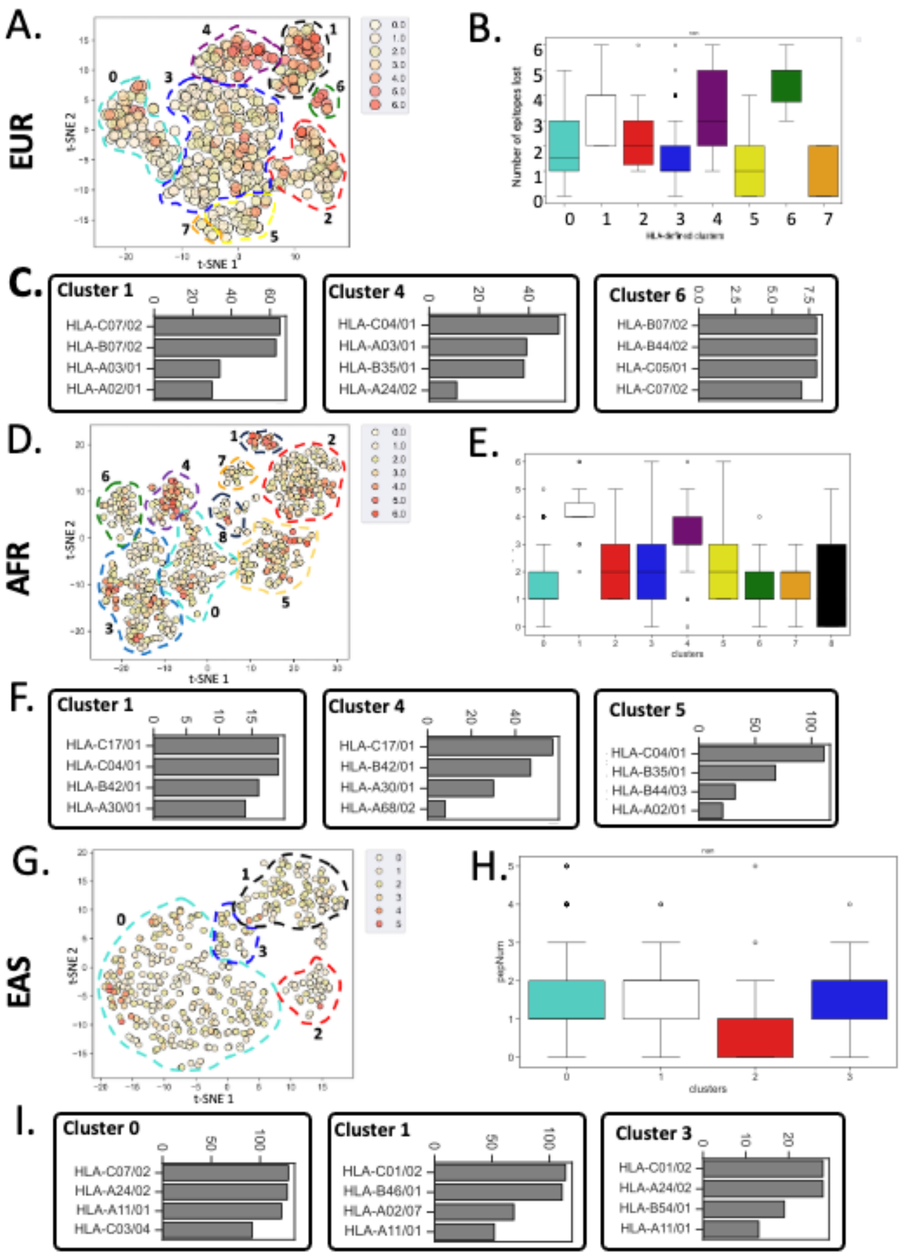

**Figure S7. predicted epitope loss across 1KGP populations.** Here, predicted epitope loss across HLA profile-defined clusters are shown across European (EUR, top), African (AFR, middle) and East Asian (EAS, bottom) populations, acquired from the 1000 Genomes Project.

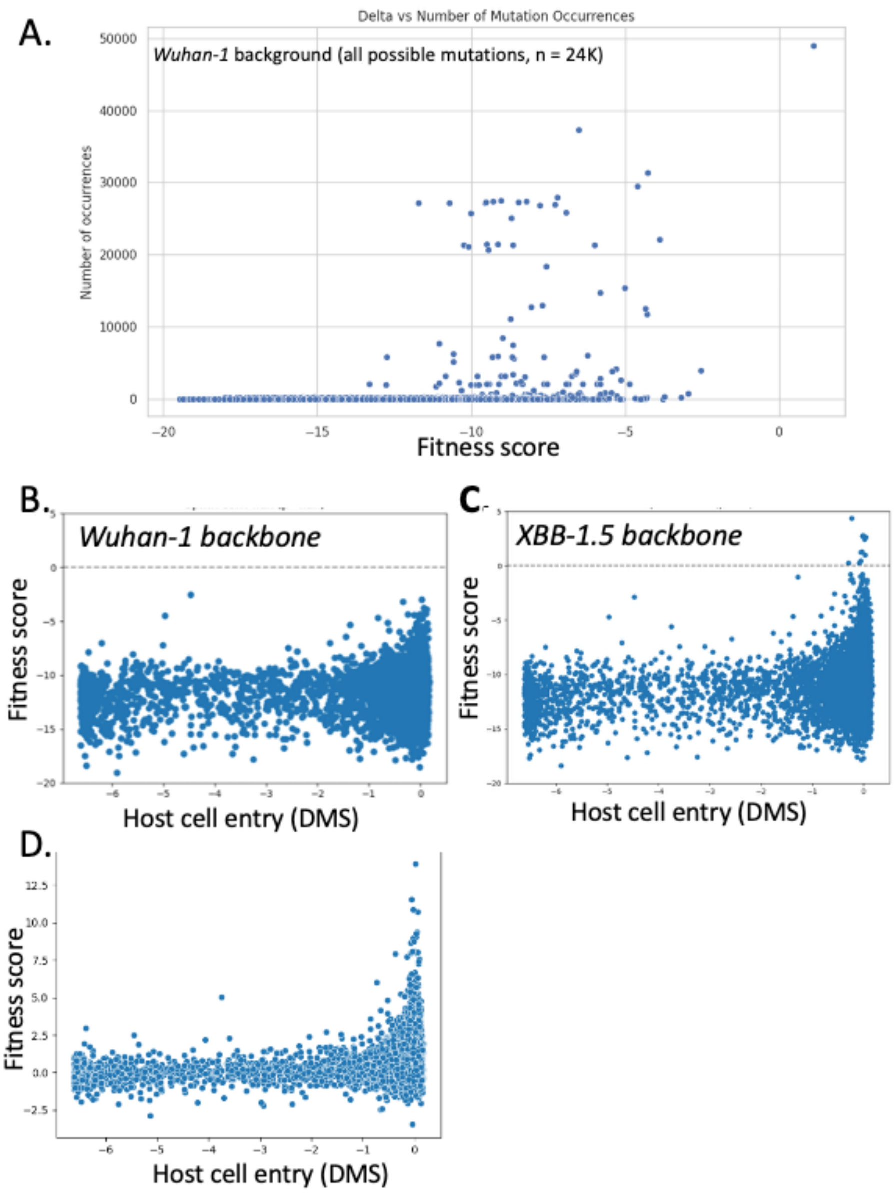

**Figure S8 Validation of PLM-based fitness estimation. A.** ESM-2-estimated fitness across all possible Spike protein mutations (n = 24K) compared to corresponding mutation occurences across a 50K GISAID sequence subsampling. **B-C**. Estimation fitness of Spike mutations assayed via DMS for Spike cell entry (Bloom et. al). The fitness of mutations were estimated using ESM-2-COV against the Wuhan-1 backbone **(B)** as well as the XBB-1.5 backbone **(C). D.** Difference in estimated fitness between mutations estimated against wuhan-1 and XBB-1.5 backbones.

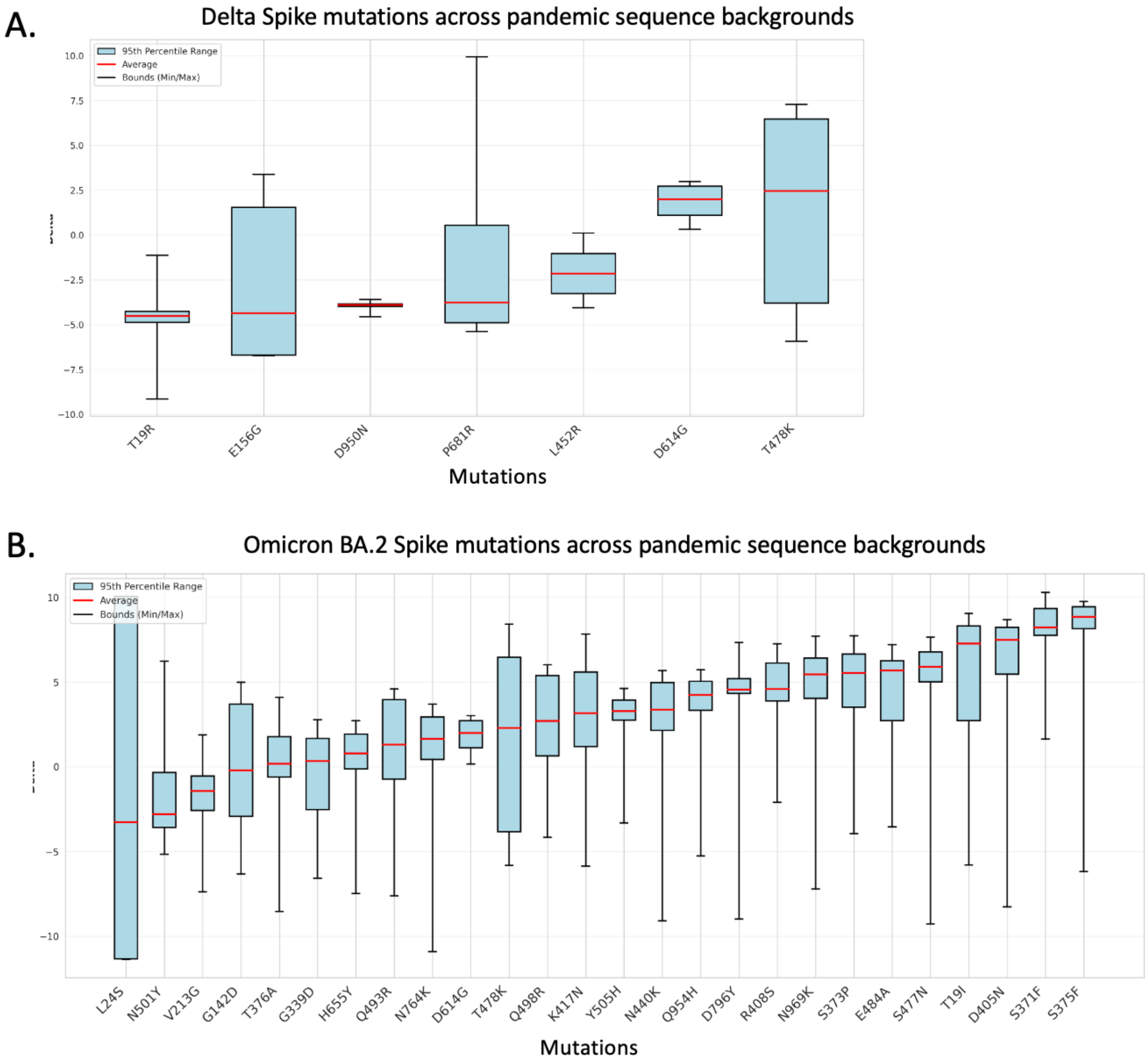

**Figure S9 Tracking the cross-pandemic fitness of mutations associated with VoCs Delta and BA.2. A**. Delta Spike mutations. **B**. BA.2 Spike mutations

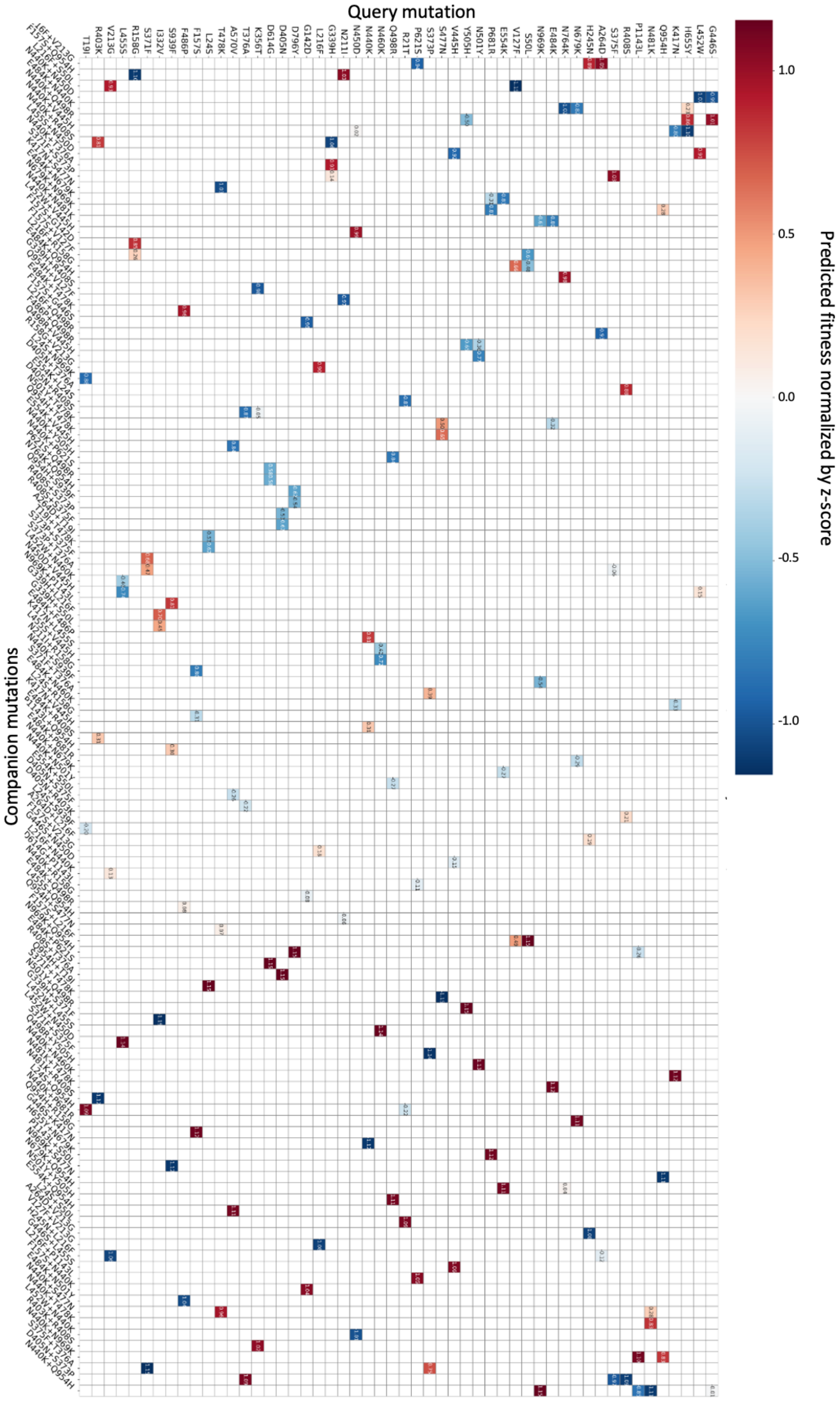
Query mutation
Companion mutations
Predicted fitness normalized by z-score
1.0
0.5
0.0
-0.5
-1.0

**Figure S10. Omicron JN.1 three-mutation interaction map.** The fitness of each Omicron JN.1 mutation was estimated in the presence of two other JN.1 mutation for possible JN.1 3-mutation combinations to create a three-mutation interaction map. Mutations are ordered by similarity in interaction profiles. Here, the y-axis corresponds the the "query mutation" (for which the fitness was assessed), while the x-axis corresponds to the "companion mutation" which was added to the backbone as a single companion mutation. Only the top 3 highest-scoring "query mutation + 2 companion mutation" groups are shown for each query mutation. Since the aim was to visualize the distribution of fitness effects of each query mutation across all companion mutations (and not to compare query mutations to each other), all fitness metrics associated with one query mutation was normalized using a z-score ((data value – average of all points)/standard deviation).

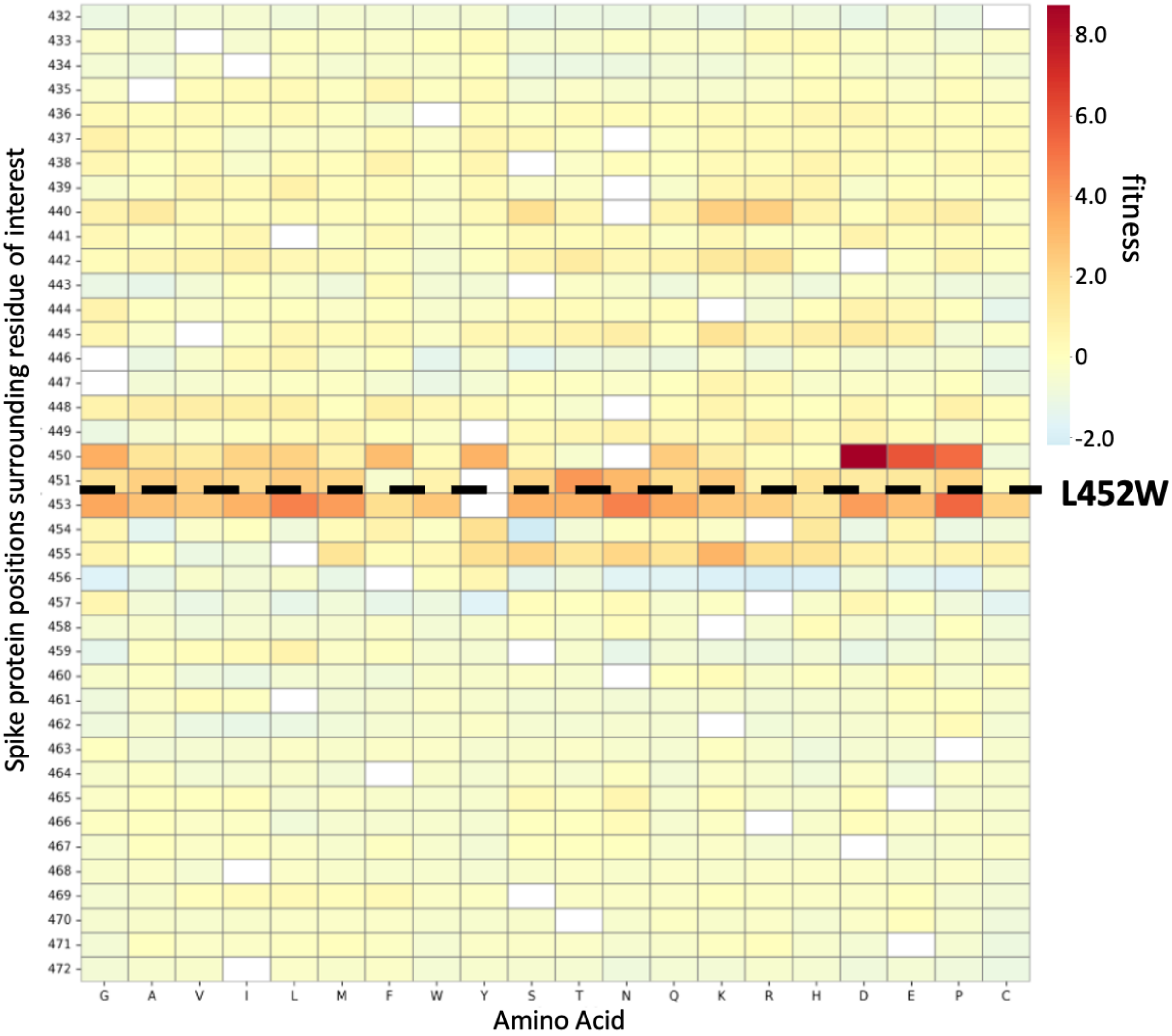

**Figure S11. Interaction map between mutation of interest (Spike L452W) and all possible amino acid mutations across a -/+ 20 position window.** The color bar corresponds to the fitness of the mutation of interest (Spike L452W) when in the presence of another mutation. All possible mutations were tested. Amino acids (x-axis) are sorted by chemical properties.

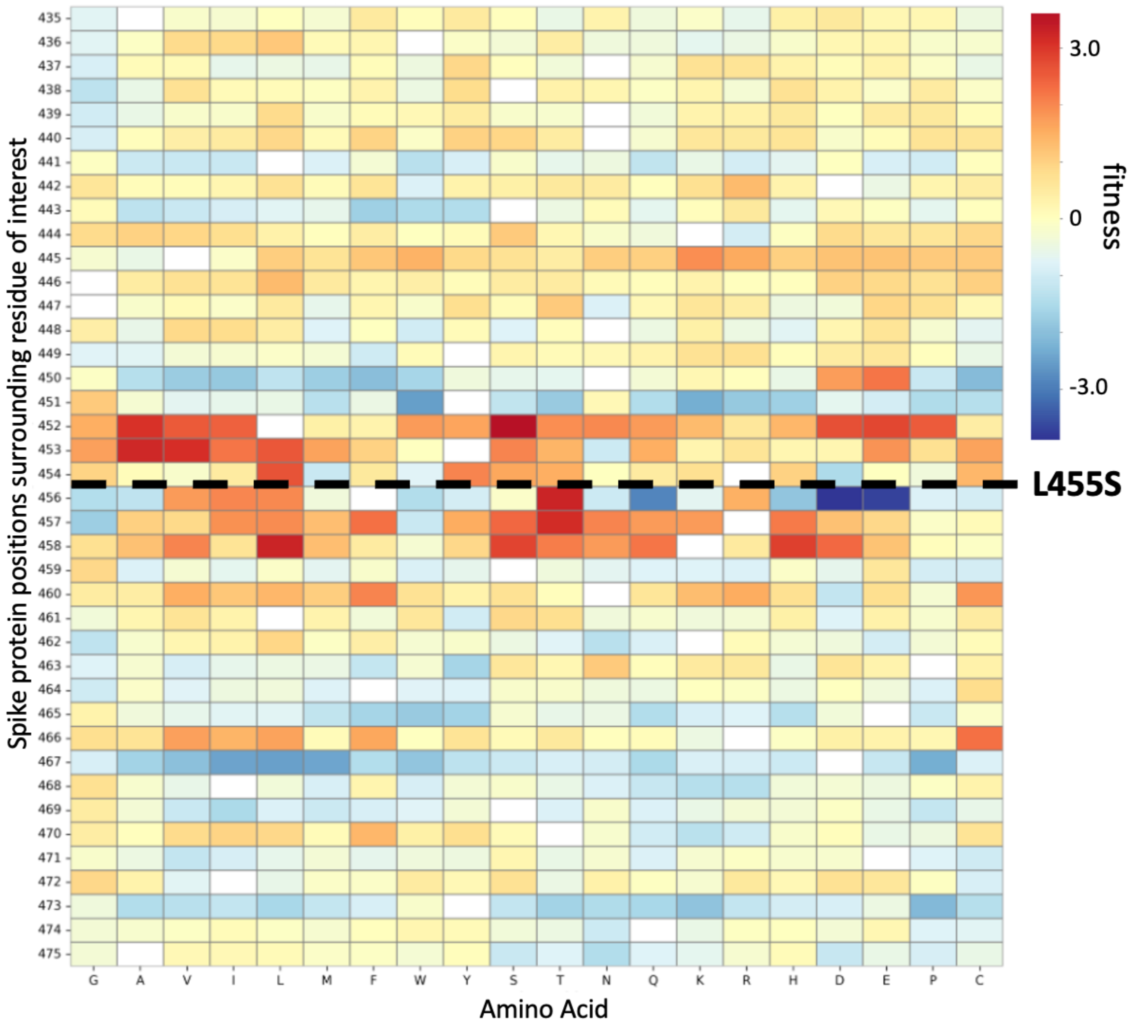

**Figure S12. Interaction map between mutation of interest (Spike L455S) and all possible amino acid mutations across a -/+ 20 position window.** The color bar corresponds to the fitness of the mutation of interest (Spike L455S) when in the presence of another mutation. All possible mutations were tested. Amino acids (x-axis) are sorted by chemical properties.

#### 5 Supplementary Table

**Table S1.** Experimentally validated SARS-CoV-2 CD8+ T Cell evasion by major variants.

| Protein | WT peptide | Mutant peptide | hla_allele | Mutation | Alpha | Delta | BA2 | JN1 | Reference |
|---|---|---|---|---|---|---|---|---|---|
| S | LLFNKVTLA | LFFNKVTLA | HLA-A02/01 | L822F | 0 | 0 | 0 | 0 | Agerer et al., Sci Immunol 2021, 'SARS-CoV-2 mutations in MHC-I-restricted |
| S | YLQPRTFLL | YFQPRTFLL | HLA-A02/01 | L270F | 0 | 0 | 0 | 0 | |
| S | NYNYLYRLF | NYNYQYRLF | HLA-A24/02 | L452Q | 0 | 0 | 0 | 0 | Deng et al., J Biol Chem 2024, 'Structural insights into immune escape at killer T cell epitope by SARS-CoV-2 |
| S | NYNYLYRLF | NYNYLFRLF | HLA-A24/02 | Y453F | 0 | 0 | 0 | 0 | |
| S | NYNYLYRLF | NYKYLYRLF | HLA-A24/02 | N450K | 0 | 0 | 0 | 1 | |
| S | YLQPRTFLL | YLQLRTFLL | HLA-A02/01 | P272L | 0 | 0 | 0 | 0 | Dolton et al., Cell 2022, 'Emergence of immune escape at dominant SARS-CoV-2 killer T cell epitope in global population' |
| S | YFPLQSYGF | YSPLRSYSF | HLA-A29/02 | F490S | 0 | 0 | 0 | 0 | Jing et al., JCI Insight 2022 T cell response to intact SARS-CoV-2 includes coronavirus cross-reactive and variant-specific components |
| N | TPSGTWLTY | KPSGTWLTY | HLA-B35/01 | T325K | 0 | 0 | 0 | 0 | Khatamzas et al., Nat Commun 2022, 'Accumulation of mutations in antibody and CD8 T cell epitopes in a B cell depleted lymphoma patient with chronic SARS-CoV-2 infection' |
| ORF1a | ALWEIQQVV | ALWEIQQFV | HLA-A02/01 | V4101F | 0 | 0 | 0 | 0 | |
| ORF1a | CTDDNALAY | CIDDNALAY | HLA-A01/01 | T4164I | 0 | 0 | 0 | 0 | |
| ORF1a | CTDDNALAYY | CIDDNALAYY | HLA-A01/01 | T4164I | 0 | 0 | 0 | 0 | |
| N | TPSGTWLTY | IPSGTWLTY | HLA-B35/01 | T332I | 0 | 0 | 0 | 0 | |
| S | FVFLVLLPLV | FVFFVLLPLV | HLA-A02/01 | L5F | 0 | 0 | 0 | 0 | Qiu et al., Front Immunol 2022, 'CD8+ T-Cell Epitope Variations Suggest a |
| S | FVFLVLLPLV | FVFLVLVPLV | HLA-A02/01 | L8V | 0 | 0 | 0 | 0 | |
| S | SPRRARSVA | SHRRARSVA | HLA-B07/02 | P681H | 1 | 0 | 1 | 0 | Swaminathan et al., Nat Commun 2022, 'Ablation of CD8+ T cell recognition of an immunodominant epitope in SARS-CoV-2 Omicron variants BA.1, BA.2 and |
| S | SPRRARSVA | SRRRARSVA | HLA-B07/02 | P681R | 0 | 1 | 0 | 1 | |
| S | YFPLQSYGF | YFPLRSYGF | HLA-A29/02 | Q493R | 0 | 0 | 1 | 1 | |
| S | YFPLQSYGF | YFPLQSYSF | HLA-A29/02 | G496S | 0 | 0 | 0 | 0 | |
| S | WMESEFRVY | WMESESRVY | HLA-A01/01 | F157S | 0 | 0 | 0 | 1 | Tian et al., Nat Immunol 2025, 'T cell immune evasion by SARS-CoV-2 JN.1 escapees targeting two cytotoxic T cell epitope hotspots' |
| S | NATNVVIKV | NATNVFIKV | HLA-B51/01 | V127F | 0 | 0 | 0 | 1 | |
| S | NATNVVIKV | NATNVFIKV | HLA-B52/01 | V127F | 0 | 0 | 0 | 1 | |
| S | NATNVVIKV | NATNVFIKV | HLA-B35/01 | V127F | 0 | 0 | 0 | 1 | |
| N | LLLLDRLNQL | LLLLDRLNKL | HLA-A02/01 | Q229K | 0 | 0 | 0 | 1 | |
| N | LLLDRLNQL | LLLDRLNKL | HLA-A02/01 | Q229K | 0 | 0 | 0 | 1 | |
| N | RLNQLESKM | RLNKLESKM | HLA-A02/01 | Q229K | 0 | 0 | 0 | 1 | |
| S | NYNYLYRLF | NYDYLYRLF | HLA-A24/02 | N450D | 0 | 0 | 0 | 1 | |
| S | NYNYLYRLF | NYNYWYRLF | HLA-A24/02 | L452W | 0 | 0 | 0 | 1 | |
| S | NYNYLYRLF | NYNYLYRSF | HLA-A24/02 | L455S | 0 | 0 | 0 | 1 | |
| S | ESIVRFPNITN | TESIVRFPNVTNL | HLA-B40/01 | I332V | 0 | 0 | 0 | 1 | |
| S | VYAWNRKRI | VYAWNRKRI | HLA-A24/02 | K356T | 0 | 0 | 0 | 1 | |
| S | RLQSLQTYV | RLQSLQIYV | HLA-A02/01 | T1006I | 0 | 0 | 0 | 0 | Wu et al., Nat Commun 2022 / JBC 2023, structural and functional studies |
| S | YLQPRTFLL | YLQLRTFLL | HLA-A02/01 | P272L | 0 | 0 | 0 | 0 | |
| S | KIADYNYKL | TIADYNYKL | HLA-A02/01 | K417T | 0 | 0 | 0 | 0 | Zhang et al., Cell Reports 2021, T-Scan profiling and structural analysis |
| S | KIADYNYKL | NIADYNYKL | HLA-A02/01 | K417N | 0 | 0 | 1 | 1 | |
| S | NYNYLYRLF | NYNYRYRLF | HLA-A24/02 | L452R | 0 | 1 | 0 | 0 | |
| S | LYNSASFSTF | LYNLASFSTF | HLA-A24/02 | S371L/F | 0 | 0 | 1 | 1 | Chung et al., 2025, Front. Immunol. |
| S | LYNSASFSTF | LYNSAPFSTF | HLA-A24/02 | S373P | 0 | 0 | 1 | 1 | |
| S | LYNSASFSTF | LYNSASFFTF | HLA-A24/02 | S375F | 0 | 0 | 1 | 1 | |
| S | LYNSASFSTF | LYNSASFSAF | HLA-A24/02 | T376A | 0 | 0 | 1 | 1 | |
| ORF1a | AANFCALILA | ADNFCALILA | HLA-A02/01 | A1708D | 1 | 0 | 0 | 0 | Xiao et al., 2022, iScience |
| ORF1a | IIWFLLLSV | TIWFLLLSV | HLA-A02/01 | I2230T | 1 | 0 | 0 | 0 | |

#### *6* Supplementary Files

Supplementary File S1 - Four-digit HLA class I alleles for the participants of the RECOVER cohort. Both the HLA calling (Tab A) as well as the associated HLA calling probability (Tab B) are provided (available on github, see data availability).

Supplementary File S2 – *Coronaviridae* sequences acquired from UniRef100 and used for the fine-tuning of ESM-2 (available on github, see data availability).

Supplementary File S3: SARS-CoV-2 sequences (acquired from GISAID) used for the viral fitness and epistasis analyzes (available on github, see data availability).

#### *7* **Supplementary References**

1. Gourraud, P.-A. *et al.* HLA Diversity in the 1000 Genomes Dataset. *PLoS ONE* **9**, e97282 (2014).

2. Stockton, J. D. *et al.* Rapid, highly accurate and cost-effective open-source simultaneous complete HLA typing and phasing of class I and II alleles using nanopore sequencing. *HLA* **96**, 163–178 (2020).

3. Dilthey, A. T. *et al.* HLA*LA—HLA typing from linearly projected graph alignments. *Bioinformatics* **35**, 4394–4396 (2019).

4. Dadonaite, B. *et al.* Spike deep mutational scanning helps predict success of SARS-CoV-2 clades. *Nature* **631**, 617–626 (2024).
